# A hypothalamus-brainstem circuit governs the prioritization of safety over essential needs

**DOI:** 10.1101/2024.09.05.611412

**Authors:** Nathalie Krauth, Lara K. Sach, Christoffer Clemmensen, Ole Kiehn

## Abstract

Animals continously adapt their behavior to balance survival and fulfilling essential needs. This balancing act involves prioritization of safety over the pursuit of other needs. However, the specific deep brain circuits that regulate safety-seeking behaviors in conjuction with motor circuits remain poorly understood. Here we identify a class of glutamatergic neurons in the lateral hypothalamic area (LHA) that target the midbrain locomotor-promoting pedunculopontine nucleus (PPN). Upon activation, this LHA-PPN pathway orchestrates context-dependent locomotion, prioritizing safety-directed movement over other essential needs such as foraging or mating. Remarkably, the neuronal activity of these circuits correlates directly with safety-seeking behavior. These circuits may respond to both intrinsic and external cues, playing a pivotal role in ensuring survival. Our findings uncover a circuit motif within the lateral hypothalamus that when recruited, prioritizes critical needs through the recruitment of an appropriate motor action.

## Introduction

The brain continuously integrates and balances internal and external cues to ensure survival by prioritizing critical needs. Navigating between the drive to fulfill essential needs and the imperative to evade imminent threats is a delicate balancing act. This requires ambulatory movement towards desired objects, amid a constant conflict with need for safety. Therefore, initiating ambulatory movement is crucial not only for approaching targets but also for shifting priorities to ensure safety when necessary.

The canonical areas directly involved in execution of ambulatory movement reside in the mesencephalic locomotor region (MLR) in the midbrain. The MLR is composed of glutamatergic neurons in the cuneiform nucleus (CnF) and the caudal pedunculopontine nucleus (PPN). CnF elicited locomotion controls both slow locomotion as well as fast escape locomotion. In contrast, caudal PPN is strictly associated with slow locomotion.^1–4^ These two regions receive synaptic inputs from many brain areas, which may facilitate the initiation of locomotion in different contexts. These contexts, include approach behaviors, such as scavenging for food, as well as movements related escape responses.^5–11^ It is imperative to delineate circuit motifs of such behaviors.

One brain area that is thought to contain circuits involved in foraging, social interaction and safety-seeking is the hypothalamus, particularly the lateral hypothalamus. These behaviors by necessity rely on the execution of ambulatory movement. Electrical stimulation of the lateral hypothalamus, as well as direct glutamate infusions can initiate locomotion in rats.^12–15^ These findings have led researchers to propose that the LHA controls appetitive locomotion funneled through the MLR.^13,16^ While an output to locomotor initiating neurons from the LHA has been described anatomically,^1,2^ it remains unknown which of the many neuronal types in the LHA^17–19^ project specifically to the PPN or CnF. It is also unclear whether LHA-mediated ambulation is recruited in specific contexts, for example linked to approach, foraging, or escape behavior. Levering cell-specific activation and inactivation experiments, along with cellular recordings combined with specific behavioral set-ups, we set out to determine this.

Here we uncover a restricted population of glutamatergic neurons in the LHA with targeted projections to the caudal locomotor promoting part of the PPN. Activation of these neurons drives ambulation in a specific, hitherto undescribed behavioral context: it overrides ongoing approach behaviors and prioritizes safety-seeking survival behavior over nutritional and social needs. The circuits can be both intrinsically activated and activated by external cues, playing an essential role in safety-seeking. Our findings uncover a circuit motif intrinsic to lateral hypothalamus that when activated prioritizes critical needs by recruiting specific brainstem locomotor pathways.

## Results

### Excitatory neurons in the LHA project to the mesencephalic locomotor region

As a first step to validate involvement of the MLR in LHA-evoked locomotor initiation, we investigated possible anatomical neuronal connections between these two regions. Previous research employing viral tracing techniques, confined to monosynaptic connections, found that locomotion driving glutamatergic neurons located in the pedunculopontine nucleus (PPN), and the cuneiform nucleus (CnF) are recipient of direct synaptic inputs from the LHA.^1,2^ However, these tracing experiments did not reveal which specific neuronal hypothalamic populations projected to the CnF or PPN. To evaluate this, we targeted excitatory (vesicular glutamate transporter2, Vglut2-positive) and inhibitory (vesicular GABA transporter, VGAT-positive) cells by injecting a AAV1-phSyn1(S)-FLEX-tdTomato-T2A-SypEGFP-WPRE, a Cre-dependent anterograde virus driving expressing cytoplasmic tdTomato and presynaptic synaptophysin-fused EGFP^20^ in the LHA of *Vglut2*^*Cre*^ or *VGAT*^*Cre*^ mice (Figure 1A). While the VGAT positive cells projected to both PPN and CnF (Extended Data Figure 1G+H), Vglut2 positive cells only project to the PPN with no projections to the CnF (Figure 1E-H, Extended Data Figure 1D+E). Vglut2 positive terminals were also found in the ventrolateral PAG close to PPN. The projections from LHA to PPN targeted the caudal part of PPN (AP -4.7 from Bregma, Figure 1E+F; Extended Data Figure 1D) which is known to be involved in promotion of locomotion.^2,4^. In contrast there are little or no projections to the rostral PPN (Extended Data Figure 1D, -4.5 mm from Bregma) which is involved in global motor arrest.^21^ These findings demonstrate a direct excitatory connection from LHA to the locomotor promoting PPN in the midbrain. To determine the distribution of input cells to the PPN in the LHA, we used a retrograde virus, retroAAV-hSyn1-chI-dloxH2BJ_mScarlet-I-NLS_EGFP(rev)-dlox-WPRE-SV40p(A) in *Vglut2*^*Cre*^ mice, which will retrogradely label neurons in the absence of cre with mScarlet, and with EGFP in cells expressing cre (Figure 1C). This approach revealed a population of cells projecting to the PPN with only a subpopulation of these being Vglut2-positive cells (Figure 1I-L). The Vglut2 positive inputs from LHA to PPN are concentrated with peak labeling along the antero-posterior LHA axis at AP -1.6 from Bregma (Extended Data Figure 1F).

**Figure 1:**
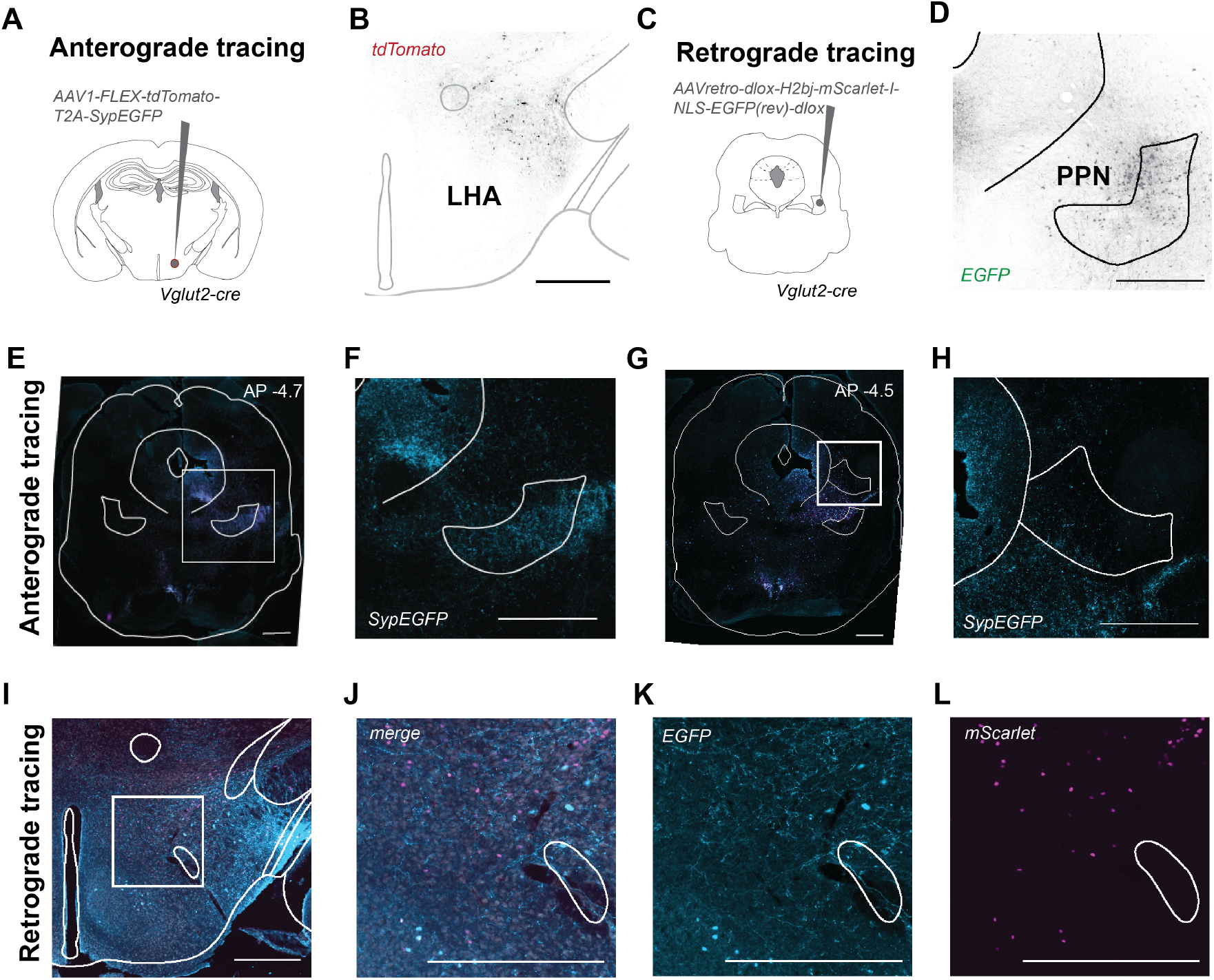
Glutamatergic projections from lateral hypothalamic area (LHA) to the brainstem locomotor area. **A)** Experimental scheme of anterograde tracing with unilateral viral injection in LHA in Vglut2^Cre^ mice. **B)** Representative image at AP -1.6 from Bregma showing labeling of Vglut2^+^ cells in LHA. **C)** Experimental scheme of retrograde viral tracing with injection in caudal PPN in Vglut2^Cre^ mice. **D)** Representative image of injection site showing Vglut2+ cells in PPN at AP -4.7 from Bregma. **E)** Representative tile images of anterogradely labeled terminal areas marked by tdTomato and SypEGFP (AP -4.7 from Bregma). **F)** Representative image of SypEGFP labeled terminals in the caudal PPN (AP -4.7 from Bregma). **G)** Representative tile images of anterogradely labeled terminal areas marked by tdTomato and SypEGFP (AP -4.5 from Bregma). **H)** Representative image of (the lack of) SypEGFP labeling in the Cuneiform nucleus (CnF; AP -4.5 from Bregma). **I)** Representative tile images of retrogradely labelled cells bodies in LHA EGFP or mScarlet (AP -1.6 from Bregma). **J)** Enlarged section of labelled cells bodies in LHA - EGFP and mScarlet merged (AP -1.6 from Bregma). **K)** Enlarged section of labeled EGFP postive cells bodies in LHA. **L)** Enlarged section of mScarlet postive cells bodies in LHA. All scale bars 500mm.

### Frequency-modulated optogenetic activation of excitatory LHA-PPN projection neurons initiates locomotion

Having demonstrated the existence of an anatomical glutamatergic lateral hypothalamic input to the caudal PPN, we investigated whether this input can drive locomotion. To specifically target the glutamatergic inputs in the LHA that project to the PPN, we administered a conditional retrograde virus expressing ChR2 in *Vglut2*^*Cre*^ mice in the caudal PPN and placed an optic fiber above the LHA (Figure 2A; Extended Data Figure 2A-B). Upon stimulation of the Vglut2 positive cell bodies in LHA with blue light (473 nm) locomotor initiation was consistently observed in an open field task.

**Figure 2:**
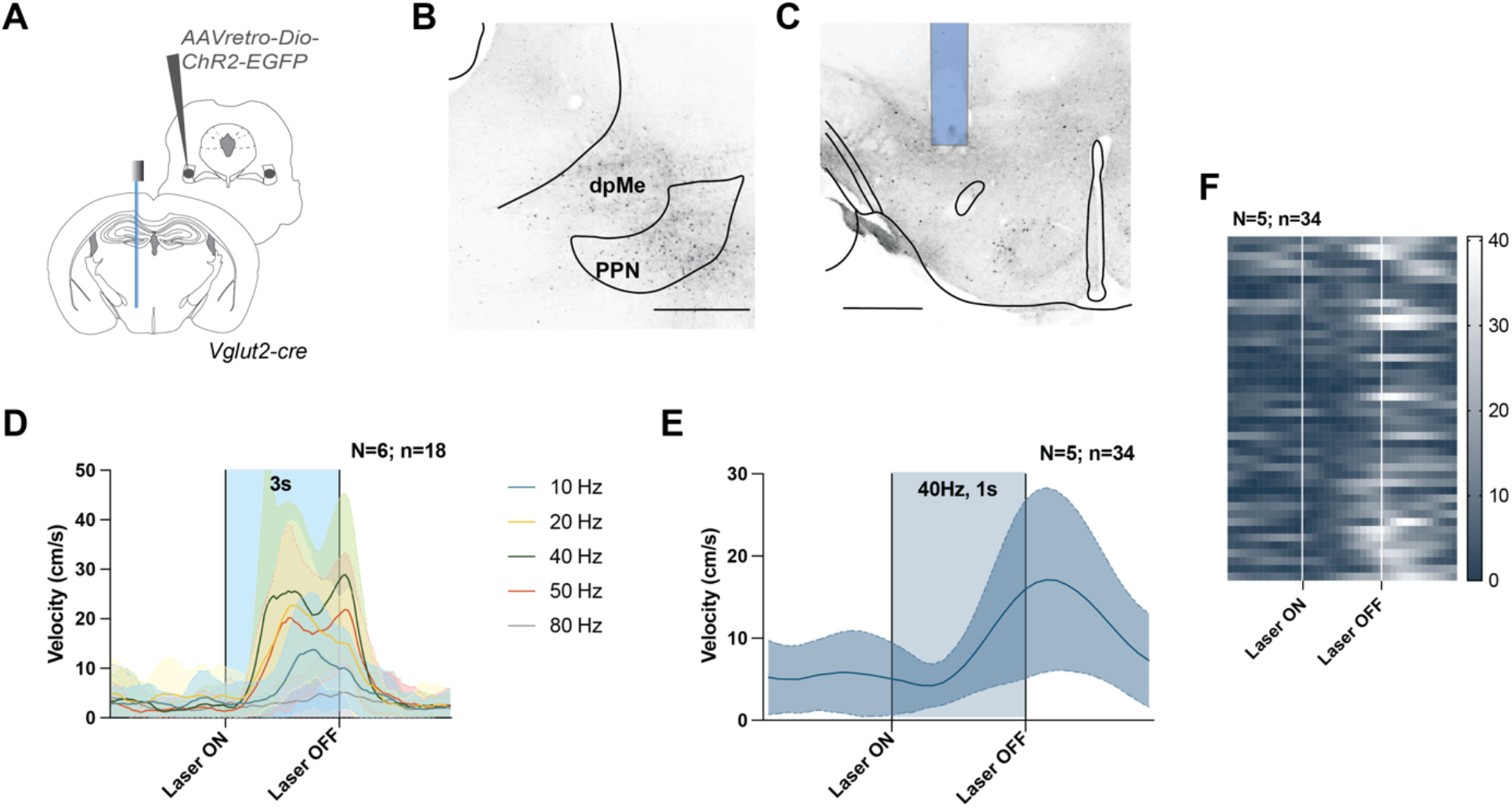
Activation of glutamatergic LHA-PPN projection neurons elicit frequency-dependent locomotion. **A)** Experimental strategy for optical activation of glutamatergic LHA-PPN projection neurons. **B)** Representative image of injection site in PPN (EGFP, AP: -4.7) showing local infection of Vglut2 positive neurons in PPN. Retrograde local labeling in deep mesencephalic nucleus (dpMe). **C)** Representative image of cell body labeling and fiber placement in LHA (EGFP, AP: -1.6, fiber placement marked with blue rectangle). **D)** Mean velocity after stimulation of glutamatergic LHA-PPN projection neurons for a duration of 3 seconds at different frequencies (N=6, n=18; 3 stimulations per mouse; mean ± SD; highlighted area represents ‘laser on’). **E)** Latency to onset of locomotion evaluated with 1 s train of pulses at 40 Hz (N=5, n=34 stimulations in total; mean ± SD; highlighted area represents ‘laser on’). **F)** Heatmap of individual experiments with 40 Hz stimulation. All scale bars: 500 mm

The velocity and latency to onset of locomotion increased as a function of stimulation frequency (10, 20, 30, 40, 50, 80 Hz; 3-second-long trains; pulse duration 20 ms) with the highest mean velocity and shortest onset latency at 40 Hz. Frequencies of 10 Hz were less efficient than 40 Hz and 80 Hz stimulation failed to evoke locomotor activity (Figure 2D). The evoked velocities are in the same range of locomotor velocities seen after direct optical stimulation of Vglut2 positive cells in the caudal PPN.^2,4^ Furthermore, 40 Hz frequency stimulation was an optimal stimulation frequency for driving the LHA-PPN projection neurons similar to the optimal frequency for direct activating Vglut2-positive neurons in PPN.^2,4^ At 40 Hz stimulation the latency to onset of locomotion from activation of glutamatergic LHA to caudal-PPN projection neurons was approximately 450 ms, reaching peak velocity at approximately 1200 ms after onset of the stimulation (Figure 2E+F). These findings show that specific activation of the glutamatergic LHA to caudal-PPN projection neurons reliably elicits forward locomotion with latencies to onset and velocities being modulated by the degree of activation but in the same range as seen by direct activation of glutamatergic neurons in caudal PPN.

### Activation of glutamatergic LHA-PPN projection neurons initiates avoidance behavior

The next question we asked was, which type of behavior is linked to the locomotion evoked by LHA-PPN projection neurons. Avoidance behavior has previously been described as a hallmark behavior linked to broad activation of glutamatergic neurons in the LHA.^22–24^ To evaluate if the movement initiated by the glutamatergic LHA-PPN projection neurons contain elements of avoidance behavior we conducted a place preference experiment. The projection neurons were targeted as in the previous experiment, with a retrograde approach from the PPN, with a fiber placed above the LHA (Figure 3A). Mice were placed in an arena with two zones of equal size (A and B) separated by walls with an opening in the middle (Figure 3B). In the first half of the experiment, simulation was coupled to zone A, while in the second half, it was coupled with zone B. This shift in stimulated zones was performed to control for any inherent side preference that the animal might exhibit. Activation of glutamatergic LHA-PPN projection neurons with blue light (473 nm) in either zone A or zone B caused active avoidance of the laser-coupled zone (Figure 3C-D, F) in contrast to control experiments stimulating with a yellow laser (593 nm) that induced no avoidance (Figure 3E-F; Extended Data Figure 3A). The mean velocity of movement was significantly increased during 473 nm laser stimulation (Figure 3G), opposed to 593 nm laser stimulation (Figure 3H). In contrast, when glutamatergic neurons in the caudal PPN were broadly activated, the mice showed no place preference (Extended Data Figure 3B-D). Together, these results suggest that activation of the specific glutamatergic LHA-PPN projection neurons causes movement to avoid a certain environment.

**Figure 3:**
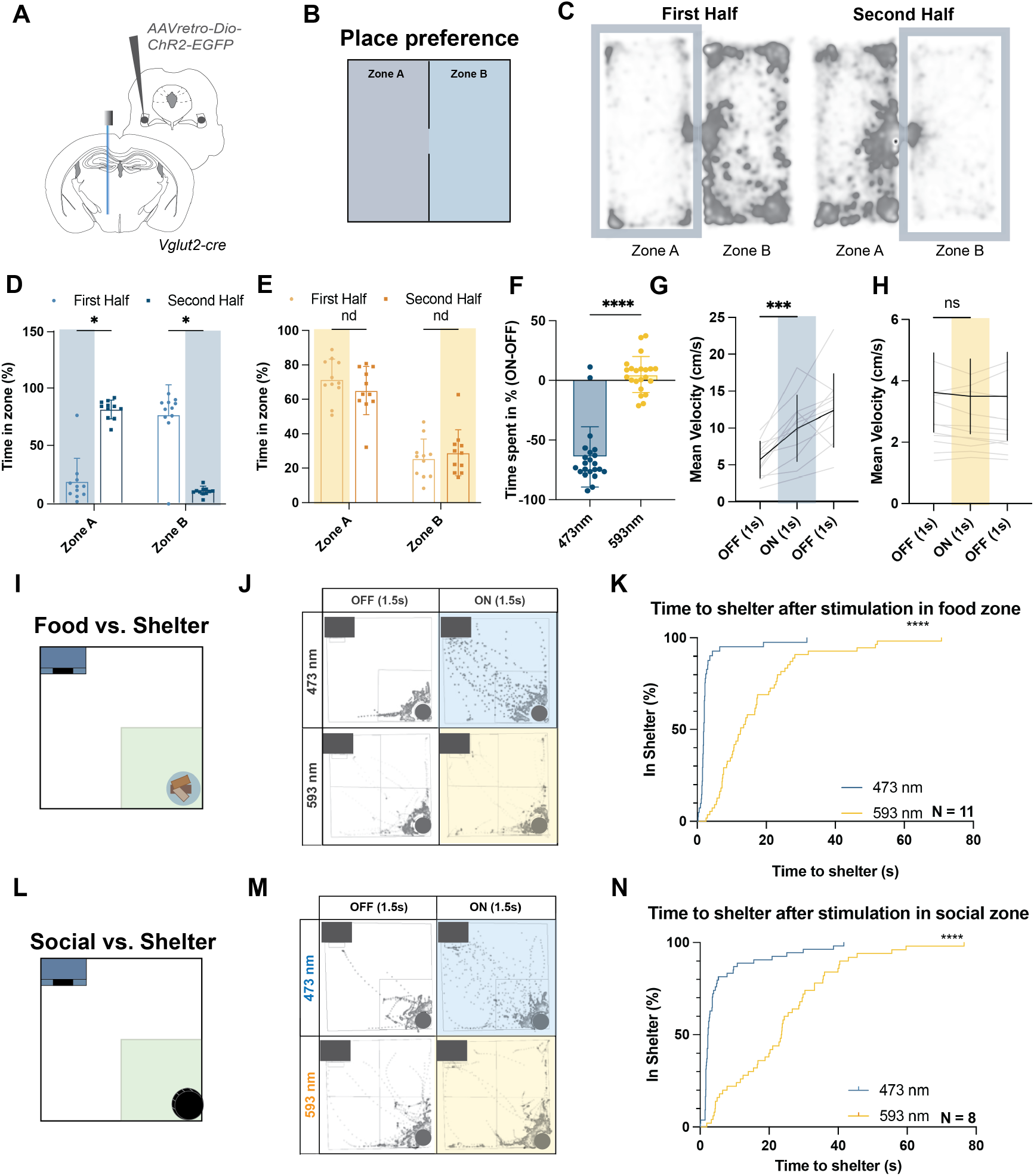
Activation of glutamatergic LHA-PPN projection neurons triggers place aversion and shelter-seeking. **A)** Experimental scheme of optogenetic targeting of glutamatergic LHA-PPN projection neurons: retroAAV-Dio-ChR2 injection in PPN (AP = 4.7 mm) with fiber placement above the LHA (AP = 1.6 mm). **B)** Arena for real-time place preference test: stimulation after 2 seconds in Zone A in the first half (0-10 mins) followed by stimulation in the second half (10-20 mins) whenever in zone B. All mice were placed in Zone A at the beginning of the trial. **C)** Heatmaps with traces of all tested mice in the first and second time period (blue frame). **D)** Cumulative time (% of total time) spent in A and B in first and second time period (Zone A: Time spent first vs. second time period (N=11; mean ± SD; paired t-test; p=0.000006); Zone B: Time spent first vs. second time period (paired t-test; p=0.0000013). Blue highlighted area represents the zone that received laser stimulation). **E)** Cumulative time (% of total time) with control laser 593 nm. (Zone A: Time spent first vs. second time period (N=11, mean ± SD, paired t-test p=0.216227); Zone B: Time spent first vs. second time period (paired t-test; p=0.455161). Yellow highlighted area represents stimulated zone. **F)** Difference score of time spent in stimulated vs non stimulated zones (N=11, n=22 halves; paired t-test p<0.0001; bar plot showing single values with mean ± SD). **G)** Mean movement velocity increases upon stimulation with 473 nm laser. Highlighted area shows laser stimulation; mean ± SD with single values in grey (N=11; paired t-test; p=0.0007). **H)** Movement velocity when mice are stimulated with 593 nm laser. Highlighted area shows laser stimulation; mean ± SD with single values in gray (N=11; paired t-test; p=0.1858). **I)** Arena for food vs. shelter experiment. Laser stimulation (1.5 sec, 40 Hz) after 1.5 seconds in food quarter. **J)** Heatmap with movement traces from all mice during pre-laser periods and laser on (1.5 sec, 40 Hz) periods. Comparison of stimulation with 473 nm activation and 593 nm control lasers. Shelter in blue. **K)** Survival plot of latency of return time to shelter after laser stimulation. Comparison of 493 nm laser and 593 nm control laser (N=11; Each ‘step’ represents one trip back to the shelter; KaplanMeier (Log-Rank) Test p<0.0001). **L)** Arena for novel mouse vs. shelter experiment. For each test a novel mouse of the opposite sex was used. Laser stimulation (1.5 sec, 40 Hz) after 1.5 seconds in the social quarter. **M)** Heatmap with mouse traces of all mice during pre-laser periods and laser on (1.5 sec, 40 Hz) periods. Comparison of stimulation with 473 nm activation and 593 nm control lasers. Shelter in blue. **N)** Survival plot of latency of return time to shelter after 493 nm activating laser and 593 nm control laser (N=8; KaplanMeier (Log-Rank) Test p<0.0001).

### Optogenetic activation of glutamatergic LHA-PPN projection neurons causes shelter directed movement in foodseeking mice

While the place preference experiments indicate an avoidance response linked to the glutamatergic LHA-PPN circuit the contextual triggers of this behavior remain to be uncovered. Avoidance driven by Vglut2 expressing cells in the LHA, has previously been linked to food aversion with mice showing decreased time spent with the food.25,26 Therefore, we set out to assess whether the avoidance response mediated by the glutamatergic LHA-PPN circuit is linked to food aversion. Hunger is a strong motivator and increases the willingness of mice to face danger to obtain food, while satiety decreases risk-taking and exploration and increases avoidance.^27–30^ With this in mind, we performed experiments in fasted animals, providing them an opportunity to seek shelter (Figure 3I). Remarkably, when glutamatergic LHA-PPN projection neurons were stimulated with blue light (473nm) in fasted mice with a high motivation to find food, the mice immediately neglected the food to move towards the shelter. This shelter-seeking behavior was temporally linked to the stimulation of glutamatergic LHA-PPN projection neurons (Figure 3J, Supplemental Video 1). In contrast the 593 nm control laser light did not induce shelter-seeking behavior (Figure 3J, Supplemental Video 1). The survival plot comparing the time from onset of all laser triggers to when the mice returned to the shelter shows a significant reduction in the return time during glutamatergic LHA-PPN projection neuron stimulation (Figure 3K). The decrease in shelter-seeking time was accompanied by increase in the locomotor velocity away from the food zone when comparing 473 nm with 593 nm laser stimulations (Extended Data Figure 3E). Since the stimulation with 593 nm did not result in direct shelter-seeking, the mice spent more time in the food zone than when stimulated with blue light and consequently accumulated more laser stimulations in this location (Extended Data Figure 3F). These experiments suggest that the behavior induced by glutamatergic LHA-PPN projection neurons represents a locomotor behavior which may lead mice to prioritize shelter-seeking over energetic needs. The behavior is instantaneous and not habituated. Since the animals may return to the food area after the pathway has been stimulated and they have rested in the shelter, the behavior itself most likely does not involve food aversion per se.

### Optogenetic activation of glutamatergic LHA-PPN projection neurons causes universal shelter-seeking movement

To determine whether the prioritization for shelter-seeking is a generalized behavior, we investigated a different situation involving a competing interest with a high drive for risk-taking. Sex hormones can modulate defense responses in both male and female mice28 and engaging in mating behavior increases risk-taking while reducing safety behaviors.31,32 Considering this, we examined whether stimulation of glutamatergic LHA-PPN projection neurons would shift the preference of conspecific investigation of the opposite sex to seeking shelter instead. Mice were placed in an arena with a shelter in one corner and a chamber containing a novel mouse of the opposite sex in the opposite corner (Figure 3L). When the experimental mouse entered the social area, the glutamatergic LHA-PPN projection neurons were stimulated. Also in this setup, the 473 nm laser stimulation caused the mice to return to the shelter with a reduced return time compared to the 593 nm control laser stimulation (Figure 3M-N, Supplemental Video 1). Additionally, stimulation with the 473 nm laser increased locomotor velocity (Extended Data Figure 3G), and given the behavioral responsiveness to the laser stimulation, the mice received fewer stimulations in the social zone with the 473 nm laser compared to the 593 nm laser (Extended Data Figure 3H). In contrast when the caudal PPN was broadly stimulated, the mice exhibited as expected increased movement but not in a specific direction of the shelter (Extended Data Figure 3I-J). Based on these results, we conclude that the observed shelter-seeking behavior extends beyond a single context and is likely a generalized behavior. Furthermore, this behavior overrides other strong needs, prioritizing safety above all. This behavior is linked to the specific pathway projecting from LHA to the PPN and is not reproduced by broad activation by glutamatergic neurons in PPN.

### Chemogenetic activation of glutamatergic LHA-PPN projection neurons shortens spontaneous shelter-seeking time and increases time spent in shelter

Mice placed in the arena containing a shelter and a food or social area perform spontaneous shelter returns and spend time in the shelter without stimulation of the LHA-PPN projection neurons (Figure 3J-K+M-N), suggesting that these behaviors are regulated by changes in LHA-PPN projection neuron activity. To illuminate this possibility, we used a chemogenetic strategy enabling a more prolonged activation of the LHA-PPN projection neurons to determine if the frequency of the spontaneous returns is affected.^33,34^ We employed a dual conditional viral strategy that utilized cre-dependent retroAAV carrying FlpO injected in the caudal PPN (Figure 4A; Extended Data Figure 4A) followed by injection of a FlpO dependent hm3D virus locally in LHA (Figure 4A; Extended Data Figure 4B). This strategy allowed us to selectively activate LHA neurons that project to the caudal PPN. A crossover study design was used for the within-mouse pairwise comparison to minimize habituation effects. Mice received intraperitoneal (i.p.) injections of 1 mg/kg Clozapine-Nitric-Oxide (CNO) or saline. Behavioral testing exposed the mice to similar arenas as in the previous optogenetic experiments (Figure 3I+L). When chemogenetically activating glutamatergic LHA-PPN projection neurons the general locomotor activity increased, as assessed by velocity in an open field arena (Figure 4B), confirming that the pathway activation enhanced overall locomotor activity. Furthermore, when mice were placed in the arena with conspecific investigation versus shelter, the mice spent less time out of the shelter when chemogenetically activated (Figure 4C). The time it took the mice to spontaneously return to the shelter was also significantly shortened in the CNO-injected group compared to the saline-injected group (Figure 4D). These data show that a long-lasting increase in glutamatergic LHA-PPN projection neuron activity promotes locomotor activity and specifically increases spontaneous shelter-seeking. This indicates that the shelter returns may be regulated by timevarying intrinsic changes in the activity of LHA-PPN projection neurons.

**Figure 4:**
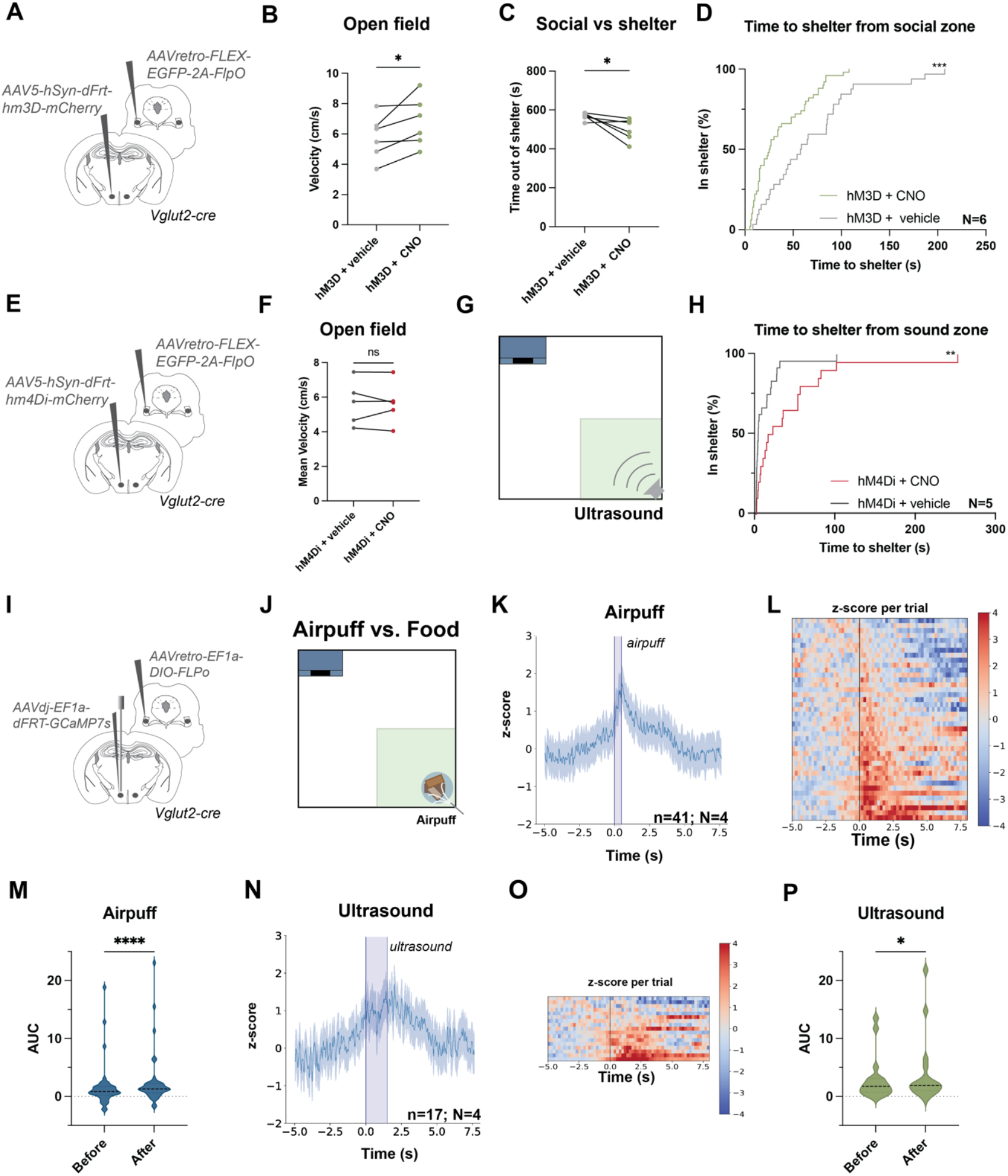
A-H. Chemogenetic activation and inhibition of glutamatergic LHA-PPN changes shelter-seeking time bidirectionally. **A)** Viral chemogenetic activation strategy of LHA-PPN projection neurons. **B)** Chemogenetic activation of LHA-PPN projection neurons leads to a slight increase in locomotor velocity in the open field. (saline versus CNO; N=6; paired t-test p=0.0464; each data point represents one animal). **C-D)** Chemogenetic activation decreases the time spent out of shelter (saline versus CNO; N=6; paired t-test p=0.0476; each data point represents one animal) (C) and decreases the return time to shelter (D). Comparison of trials after CNO and saline injections (N=6; Kaplan-Meier-Log-Rank Test p=0.0003). **E)** Chemogenetic dual viral strategy with bilateral injections of AAV-retro-FLEX-EGFP-2A-FlpO in PPN and AAV5-hSyn-dFrt-hm4Di in LHA. **F)** Chemogenetic inhibition does not affect movement velocity in the open field (saline vs CNO; N=5; paired t-test p=0.9015; each data point represents one animal). **G)** Experimental arena for shelter-seeking after ultrasound exposure. **H)** Return time to shelter after chemogenetic inhibition (N=5; Kaplan-Meier-Log-Rank Test p=0.0091). **I-P. Glutamatergic LHA-PPN neuron activity correlates with shelter-seeking. I)** Viral strategy and fiber placement to target glutamatergic LHA-PPN projection neurons for calcium imaging with fiber photometry. Dual viral strategy with injections of AAV-retro-EF1a-DIO-FlpO in PPN and AAVdj-EF1a-dFRT-GcaMP7s in LHA, with fiber placed in LHA. Experimental arena to test competing interest of food versus airpuff with a shelter present. **K)** Z-scored averaged activity aligned to onset of airpuff (Airpuff duration 500 ms; total time in seconds with total time 5 seconds before and 8 seconds after onset; Mean +/SD, N=4, n=41 trials). **L)** Heatmap of z-scores of all single airpuffs aligned to onset of airpuff at black line. **M)** Area under the curve measurement for 1 second before and after airpuff onset shows a clear increase in calcium signal after the puff (Wilcoxon matched pairs signed rank test p<0.0001). **N)** Z-scored averaged activity aligned to onset of ultrasound (ultrasound duration 1500 ms; pre-onset-time is 5 s post-onset-time is 8 s; Mean +/SD; N=4, n=17 trials). **O)** Heatmap plot of z-scores for every trial. **P)** Area under the curve measurement from 2 seconds before to 2 s after ultrasound onset (Wilcoxon matched pairs signed rank test p=0.0067).

### Chemogenetic inhibition of glutamatergic LHA-PPN projection neurons has no effect on general locomotor behavior but delays cue-dependent shelter-seeking

We next set out to evaluate if dampening the activity of the glutamatergic LHA-PPN projection neurons would affect the shelter-seeking. To investigate this, we performed specific chemogenetic inactivation of the pathway using inhibitory DREADDS (hm4Di).^33,34^ We used a dual conditional viral strategy that utilized cre-dependent retroAAV carrying FlpO injected in caudal PPN (Figure 4E; Extended Data Figure 4C) followed by injection of a FlpO dependent hm4Di virus locally in LHA (Extended Data Figure 4D). Consistent with the activation approach, we used the same crossover study design to avoid habituation. First, we tested if chemogenetic inactivation would affect general locomotor activity. Unlike the effects observed with chemogenetic activation of the pathway, there was no difference in velocity or distance moved after CNO injection compared to saline controls. This suggests that glutamatergic LHA-PPN projection neurons are not involved in regular self-paced locomotion in the open field (Figure 4F). To investigate the contribution of the pathway to shelter-seeking, we used a condition that reliably evokes shelter-seeking: a short exposure (1s) to ultrasound (20kHz). Interestingly, it has been shown that ultrasound activates neural activity in the lateral hypothalamus, as demonstrated by c-Fos staining.^35^ The ultrasound was placed in the opposite corner of the shelter and triggered when the mice entered this corner (Figure 4G). When we compared the behavior of saline-injected mice to CNO-injected mice, we observed a clear delay in ultrasound-induced shelter-seeking after CNO-injection (Figure 4H). This suggests that the cue evoked shelter-seeking behavior is reduced by specifically dampening glutamatergic LHA-PPN projection neuron activity, and that this effect is not due to a general impairment of the animal’s locomotor capabilities.

### Glutamatergic LHA-PPN projection neurons show neural activity during cue dependent shelter-seeking

The question that arises after the activation and inactivation experiments is whether LHA-PPN projection neuron activity correlates with the observed behavior. For this, we recorded the LHA-PPN projection neuron activity using fiber photometry. We used a two-step viral approach, injecting a cre-dependent retroAAV carrying FlpO into the PPN and a FlpO-dependent GCaMP virus into the LHA. The optical fiber for photometry was positioned above the LHA site where GCaMP was injected (Figure 4I, Extended Data Figure 4E). The mice were then placed in a competing interest arena to test the effect of exposing mice to an airpuff, while having the option to seek shelter (Figure 4J). Notably, when exposed to the airpuff mice consistently sought shelter. Concurrently, we observed a distinct increase in neural activity in the glutamatergic LHA-PPN projection neurons at the onset of the airpuff-induced shelter-seeking (Figures 4K-M). In contrast, when mice were exposed to the sound of the airpuff without the airpuff, there was no shelter seeking and no increase in the calcium response (Extended Data Figure 4F-G). A similar increase in glutamatergic LHAPPN projection neuron activity was observed in a shelter versus food and ultrasound arena, although less pronounced than after the airpuff (Figure 4N-P). This response difference corresponded to a more delayed and less pronounced shelter seeking after the ultrasound cue as compared to the airpuff cue, in the face of a strong (food) attractant. Together these experiments show that glutamatergic LHA-PPN projection neuron activity is strongly correlated with shelter-seeking behaviors.

### Glutamatergic LHA-PPN projection neurons have collaterals to VTA and LC but not LHb

To determine if other brain areas are involved in the prioritization of shelter-seeking, we investigated if glutamatergic LHA-PPN projection neurons had collateral projections to other brain areas. We focused on known projection areas from the Vglut2 positive neurons in LHA such as the lateral habenula (LHb),^24^ the ventral tegmental area (VTA),^23,36^ and the locus coeruleus (LC).^37,38^ To assess the collateral projections, we used a dual-color retrograde tracing strategy with cre-dependent retroAAVs injected into the PPN and either the VTA, LHb or LC of Vglut2Cre mice (Figure 5A). After allowing for virus expression, we counted single or dually retrogradely labeled cell bodies along the anterior-posterior (AP)-axis of LHA (Figure 5B). When examining the viral expression in VTA and PPN projection neurons we observed that LHA-VTA projection neurons are more prevalent in the posterior part of the LHA, while the LHA-PPN projection neurons are spread along the entire AP-axis. Notably, about 25% of the counted cells had projections to both areas particularly around the coordinates -1.6 and -1.8, the hot spot for inducing locomotion. The LHA-LHb projection neurons are mostly expressed in the anterior part of the LHA, with almost no overlap with LHAPPN projection neuron cell bodies (2.6%). The location of the LHA-LC projection neurons along the LHA AP-axis was similar to those of the LHA-PPN projection neurons with an overlap of about 14% LHA-LC and LHA-PPN projection neurons (Figure 5N). These data suggest the possibility that projections to the VTA and LC may be coactivated when stimulating the glutamatergic LHA-PPN cell bodies, potentially contributing to the shelter-seeking behavior.

**Figure 5:**
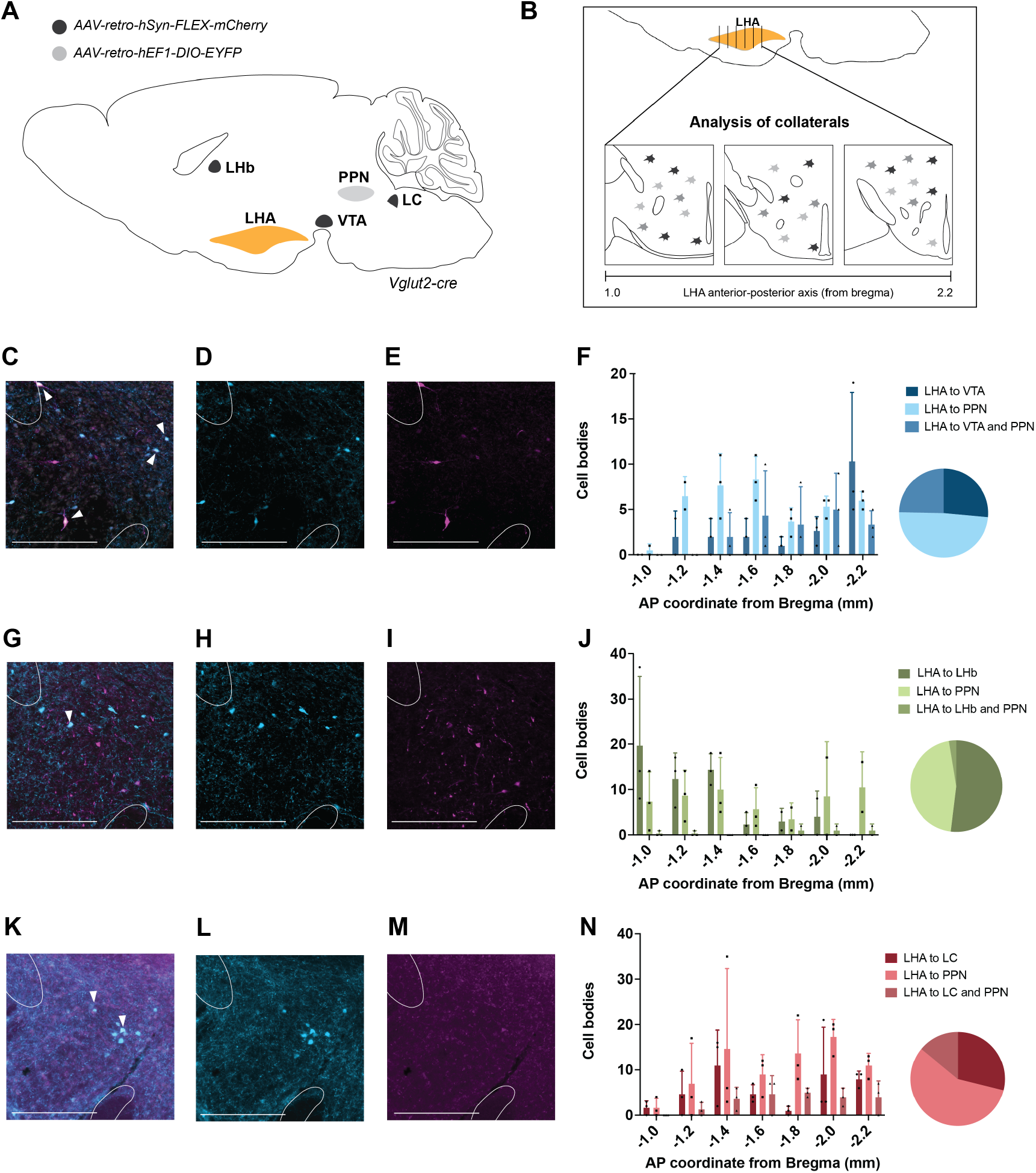
Glutamatergic LHA-PPN projection neurons have collaterals to locus coeruleus and ventral tegmental area. **A)** Dual retrograde viral injection in locus coeruleus (LC), lateral habenula (LHb) or ventral tegmental area (VTA) combined with retrograde injection in caudal PPN to label Vglut2 positive projection neurons in LHA. **B)** Coronal brain slices of the LHA along the anterior-posterior axis. **C-E)** Representative images of VTA and PPN retrogradely labeled cells bodies in LHA at AP -1.6. **C)** Overlay of both channels with arrows indicating double-labeled cell bodies. **D)** Cell bodies projecting to PPN. **E)** Cell bodies projecting to VTA (scale bars 500 mm). **F)** Count of cell bodies retrogradely labeled cells from PPN and VTA. Means +/SD; N=3. Pie chart represents the percentage of LHA cell bodies projecting to PPN (48.86%), VTA (26.48%) or both (24.66%). **G-I)** Representative images of LHb and PPN retrogradely labeled cell bodies in LHA at AP -1.6. LHA showing separate channels and overlay of both channels with arrows indicating double labeled cell bodies (scale bars 500 um). **J)** Count of cell bodies retrogradely labeled cells from PPN and LHb. Means +/-SD; N=3. Pie chart represents the percentage of LHA cell bodies projecting to PPN (45.45%), LHb (51.95%) or both (2.60%).**K-M)** Representative images of LHb and PPN retrogradely labeled cell bodies in LHA at AP -1.6 showing separate channels and overlay of both channels with arrows indicating double labeled cell bodies (scale bars 500 um). **N)** Count of cell bodies retrogradely labeled cells from PPN and LC. Means with +/-SD; N=3. Pie chart represents the percentage of LHA cell bodies projecting to PPN (57.03%), LC (28.91% or both (14.06%).

### Inhibition of Vglut2^+^-VTA neurons does not affect glutamatergic LHA-PPN projection neuron driven shelter-seeking behavior

Recent research has shown that glutamatergic LHA-VTA projection neurons are involved in food-seeking behavior. This effect is specifically mediated by cells that project onto a VTA population of neurons that express Vglut2.^39,40^ To evaluate the potential contribution of co-activation of LHAVTA projection neurons to the observed behavior, we conducted an experiment where the VTA-Vglut2 population was chemogenetically inhibited during glutamatergic LHAPPN optogenetic stimulation (Extended Data Figure 6C). Interestingly inhibition of the VTA-Vglut2 population did not alter shelter-seeking time or movement upon laser stimulation (Extended Data Figure 6C-G), suggesting that functional effect of the glutamatergic LHA-PPN subpopulation is distinct from the LHA-VTA projections onto glutamatergic VTA neurons.

**Figure 6:**
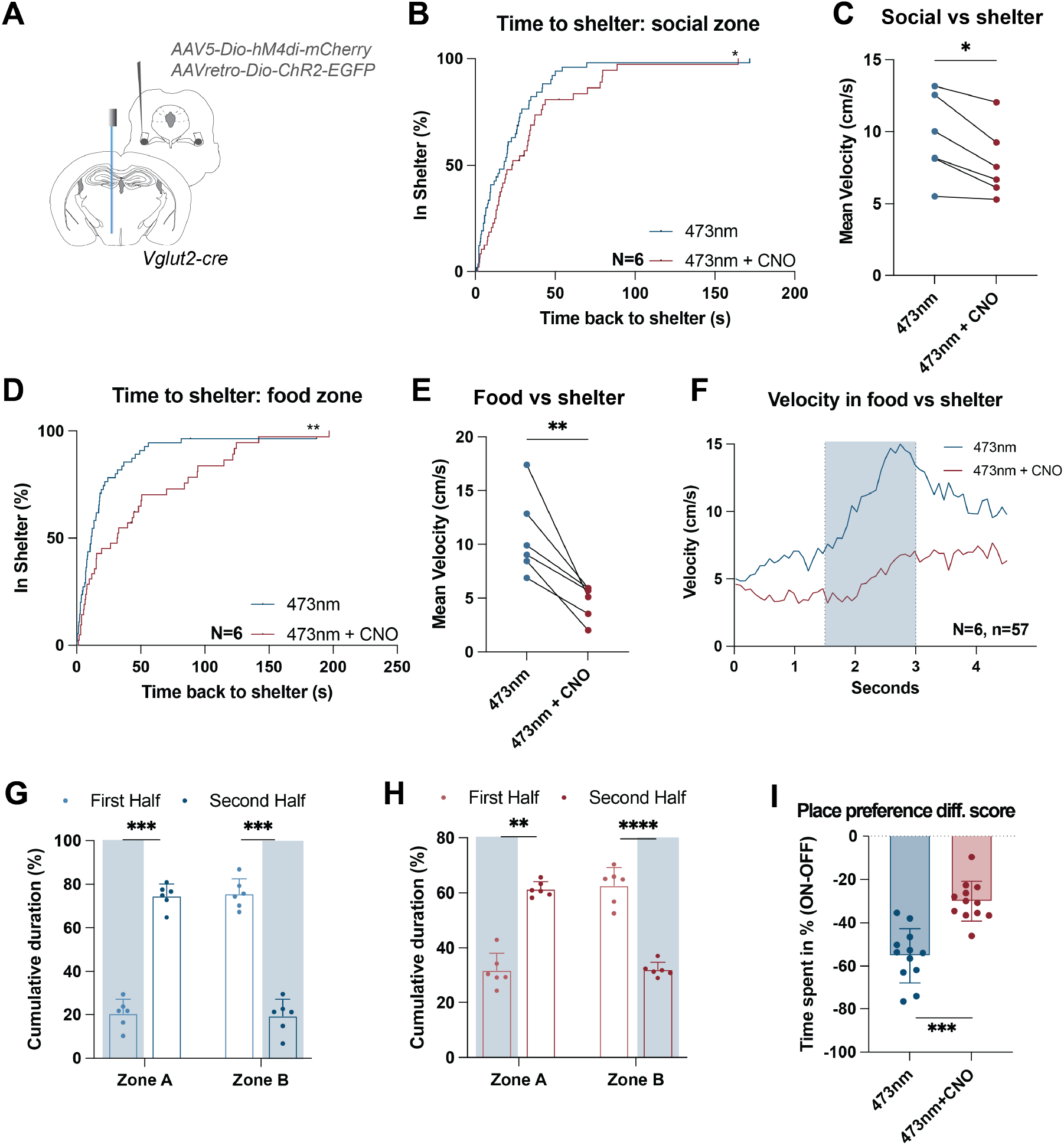
LHA-PPN projection neuron evoked shelter-seeking and place aversion is dampened by inhibition of PPN. **A)** Experimental strategy with bilateral injection of the AAV-hM4di in PPN and retrograde expression of ChR2 in glutamatergic LHA-PPN projection neurons. **B)** The return latencies are prolonged after optical stimulation of glutamatergic LHA -PPN projection neurons in the social zone after PPN inhibition. Comparison of saline injected and CNO injected mice (N=6; KaplanMeier (Log-Rank) Test p=0.0437). **C)** Comparison of mean velocity during 1.5 second long laser stimulations shows reduced locomotor velocity when PPN is chemogenetically inhibited (CNO vs. saline injected; N=6; paired t-test p=0.0100; each data point represents one animal). **D)** The return latencies are prolonged after optical stimulation of glutamatergic LHA -PPN projection neurons in the food zone after PPN inhibition. Comparison CNO and saline treated mice (N=6; KaplanMeier (Log-Rank) Test p=0.0026). **E)** Comparison of mean velocity during laser stimulation shows reduced locomotor velocity when PPN is chemogenetically inhibited (CNO vs. saline injected; N=6; paired t-test p=0.0076). **F)** Mean velocity of 1.5 s long locomotion triggers in food vs shelter arena. Blue curve represents mean of velocity data points measured every 0.08 seconds in vehicle treated group, while red represents the CNO treated groups (n=6, 57 stimulations in total; highlighted area represents ‘laser on’). **G)** Place preference experiment with saline treated groups. Graph shows time spent as % of total time. (Zone A: Time spent first vs. second half (N=6; paired t-test; p=0.0002); Zone B: Time spent first vs. second half (paired t-test; p=0.0001). Blue highlighted area represents laser stimulation (N=7; bar plot of mean ± SD). **H)** Place preference experiment after PPN was inhibited with CNO. Time spent in zones in the stimulated area upon stimulation with a 473nm laser. Graph shows time spent as % of total time. (Zone A: Time spent first vs. second half (N=6; paired t-test; p=0.0020); Zone B: Time spent first vs. second half (N=6; paired t-test; p<0.0001). Blue highlighted area represents laser stimulation (N=7; bar plot of mean ± SD). **I)** Difference score of time spent in stimulated vs non stimulated zones (N=6, n=12 halves; paired t-test p= 0.0003; bar plot of single values with mean± SD).

### Inhibition of Vglut2^+^-PPN neurons reduces glutamatergic LHA-PPN projection neuron driven shelter-seeking behavior

Considering the projections of the anatomical branching from the LHA, our behavioral findings alone do not establish a definitive role for the PPN in the observed shelter-seeking behavior. To evaluate the specific role of glutamatergic PPN neurons in these behaviors, we conducted experiments with ChR2 expressed in glutamatergic LHA-PPN projection neurons while simultaneously chemogenetically inactivating glutamatergic neurons in the caudal PPN (Figure 6A, Extended Data Figure 6A-B). To avoid confounding factors due to habituation, a crossover study was chosen for the chemogenetic experiments. Inhibition of caudal glutamatergic PPN neurons with CNO significantly delayed glutamatergic LHA-PPN projection neuron-induced shelter-seeking time in the social vs. shelter experiment when compared to saline injected mice (Figure 6B). Moreover, the mean velocity induced by the 473nm laser stimulation was attenuated when comparing the CNO and saline groups (Figure 6C), showing that PPN activation was reduced. These findings demonstrate that the caudal glutamatergic PPN neurons play an essential role in driving the movement in shelter-seeking behavior, when mice choose between conspecific investigation and safety. Importantly, this role is universal for shelter-seeking. Thus, inhibition of caudal glutamatergic PPN neurons also delayed shelter-seeking time in a food versus shelter setting (Figure 6D) and reduced the mean velocity during 473 nm stimulation (Figure 6E+F). From these experiments in both the food vs. shelter and social vs. shelter settings, we conclude that PPN-directed locomotion plays a vital role in glutamatergic LHA-PPN induced shelter-seeking behavior.

### Inhibition of Vglut2^+^-PPN neurons during stimulation of glutamatergic LHA-PPN projection neurons attenuates velocity and weakens avoidance

Finally, we investigated if the avoidance behavior observed in the place preference arena was affected by chemogenetic inhibition of the glutamatergic neurons in the caudal PPN during optogenetic stimulation of glutamatergic LHA-PPN projection neurons. The place preference difference score, i.e., the difference in time spent in the stimulated vs non stimulated zones, was significantly reduced in response to the chemogenetic inhibition (Figure 6G-I). These findings show that the glutamatergic LHA-PPN projection induced place preference, is mediated through caudal glutamatergic PPN-driven locomotion, aligning with observations made during shelter-seeking behavior.

## Discussion

Our study uncovers a direct excitatory circuit from the LHA to the locomotor promoting PPN. Activation of this pathway leads to a context-dependent expression of locomotion with prioritization of safety-directed locomotion that overrides other – hypothalamic-controlled – essential needs such as foraging or mating. Our results thus reveal a circuit motif in the lateral hypothalamus that when recruited prioritizes competing needs through recruiting of an appropriate motor action. Together, our results demonstrate the LHA plays a central role not only in fundamental survival behaviors but also in dynamically adjusting these behaviors based on the current environmental context and potential threats.

Anatomical tracing studies have shown that the MLR receives direct input from the hypothalamus.^1,2,41–43^ While these studies identified the projections by employing either conventional tracing or transsynaptic virus tracing, they did not identify the input cell types in LHA. Our experiments reveal that Vglut2-LHA neurons specifically provide input to the caudal PPN, the MLR region responsible for eliciting slow to medium fast forward locomotion.^2^ In contrast, no Vglut2-LHA inputs were observed in the cuneiform nucleus (CnF), the other MLR subregion that encodes fast speed locomotion, essential for escape-like behaviors. These anatomical connections align with the findings from the optogenetic activation experiments, which demonstrate that glutamatergic LHA-PPN projection neurons reliably activate locomotion in a frequency-dependent manner with an optimal frequency around 40 Hz, corresponding to the optimal stimulation frequency of glutamatergic neurons directly in the caudal PPN.^2,4^ Moreover, the evoked locomotion exhibited a maximum velocity of approximately 40 cm/s, which is consistent to what is observed by direct stimulation of glutamatergic neurons in the caudal PPN.^2^ Collectively, these observations demonstrate that excitatory LHA neurons can access the MLR directly through selective targeting of the caudal PPN. This locomotor activity is not dependent on CnF activity and is in a locomotor range typically evoked from direct PPN stimulation. This notion is also supported by the finding that the LHA-PPN projection neuron evoked movement is severely attenuated when caudal glutamatergic PPN neurons are inactivated. The significance of this finding is that it directly links LHA-mediated locomotion to the caudal PPN and does not imply an activation of neurons in the medulla that eventually act on the spinal locomotor circuits.^2,8,16,44^

Previous experiments have shown that activation of Vglut2 expressing LHA neurons reduces food seeking and that this behavior reflects food aversion.^18,24^ Though no previous studies show projection-specificity to the PPN, our findings seem to support this conclusion at first glance. However, when we presented hungry mice with a choice between eating and seeking shelter for safety, activation of LHA-PPN projection neurons redirected the animals’ focus from interaction with the food to shelter-seeking —a distinct and competing survival behavior. Furthermore, when mice were presented with the option to investigate a conspecific instead of food, they also consistently exhibited shelter-seeking behavior in response to stimulation of LHA-PPN projection neurons. Accordingly, the shelter-seeking behavior governed by the LHA-PPN projection neurons is a generalized behavior that prioritizes safety over other strong survival behaviors and is not specifically related to food avoidance. Agreeing with this observation, the prioritization of safety over food is also evoked in fasted animals. The shelter-seeking responses are followed by a distinct and timed-locked increase in LHA-PPN projection neuron activity suggesting that their activation is related to the prioritization of safety itself rather than the specific cue triggering the shelter-seeking behavior.

Importantly, the behavioral response evoked by the LHAPPN projection neurons does not align with a typical escape response triggered by imminent threat from predators or intense fear. These responses mediated by visual stimuli activate the superior colliculi and reach the MLR directly or indirectly.^5,6,8,45^ These responses result in rapid escape or the activation of amygdala-mediated fear responses, which act on the dorsal PAG, inducing fast running and jumping that is characteristic of flight.^46^ In contrast, the characteristics of shelter-seeking behavior are more akin to a slower PPN-evoked locomotion. However, while chemogenetic activation of LHA-PPN projection neurons increased the speed of locomotion, inactivation did not, suggesting that the evoked locomotion differs from regular self-paced locomotion observed during explorative behavior.^2^ Rather LHA-PPN projection neurons are specifically recruited to control safety-seeking behavior, in which the time to return to the shelter was specifically reduced by activation of LHAPPN projection neurons and increased by their inactivation. These findings suggest that the lateral hypothalamic circuit for safety-seeking causes locomotion upon reaching a specific threshold, thus being a crucial survival mechanism. It may be controlled by both internal homeostatic signaling and external cues. Consequently, the LHA-PPN projection pathway represents a hitherto undisclosed mechanism through which the MLR (PPN/CnF) is activated by upstream brain regions, including the basal ganglia, cortex, or superior colliculus.

These findings opened avenues for further investigation into the influence of internal states on behavior. Therefore, we were interested in determining the potential interactions between the excitatory LHA-PPN projection neurons and other brain regions that are known to receive output from LHA, namely the LHb, VTA, LC, and PAG. Projections from the LHA to LHb have been demonstrated to facilitate active avoidance, aversion, and escape from a looming stimulus by releasing glutamate in the lateral habenula.^24,47^ Despite the apparent similarities between the LHA-LHb and LHA-PPN elicited behaviors, our anatomical findings refuted this assumption. There is no overlap between the LHA-LHb and LHA-PPN projection neurons in the LHA, indicating distinct pathways for these projections. This finding is compatible with the evidence presented by Rossi et al. demonstrating that the LHb receives inputs primarily from the anterior LHA, while PPN-projecting cell bodies are situated more posteriorly.^36^ As another collateral area we found the locus coeruleus (LC), which is known to send and receive inputs to the LHA.^48–50^ Anatomically, we observed an overlap of cell bodies projecting to both the LC and the PPN, suggesting a possible co-facilitation. Previous studies have demonstrated that activation of LHA-LC projecting neurons induces freezing responses,^38^ which is not compatible with our findings. However, co-activation of the LC may contribute to increased alertness and readiness to seek safety, given its role as a noradrenergic hub.^51,52^ We also identified the presence of collateral projections from the LHA to the PPN and the VTA. Extensive research has been conducted on the projections from the LHA to the VTA. These inputs include both glutamatergic and GABAergic projections to dopamine (DA) and glutamate VTA neurons.^23,53^ GABAergic inputs have been demonstrated to increase feeding behavior, while glutamatergic inputs have been shown to cause aversion, avoidance, and decreased food intake.^23,36,39,40,53,54^ It is therefore possible that some of the observed behavioral valence is mediated by VTA collaterals. However, when we inactivated Vglut2 cells in the VTA upon stimulation of LHA-PPN projection neurons, we did not observe a change in the evoked behavior, suggesting that Vglut2 cells in VTA do not play a significant role for the herein uncovered behavioral circuit. Finally, one might consider the involvement of the PAG in the observed behaviors. However, a close examination of the escape behavior observed in Li et al. 2018 reveals that the mice exhibited jumping and escaping at very high speeds.^46^ This is in stark contrast to the slower, safety-seeking behavior that we observed. It is therefore unlikely that these projections are involved the shelter-seeking behavior. Altogether, these findings indicate that the direct projection from LHA to PPN is the primary driver of the shelter-seeking behavior. This provides further evidence of the significance of this direct circuit in movement control and is corroborated by the observation that safety-seeking locomotion and place avoidance are significantly attenuated when glutamatergic caudal PPN neurons are inactivated. Exploring the broader implications of this prioritization mechanism provides an opportunity to gain a deeper understanding of the brain’s control of survival behaviors. Importantly, as this neuronal circuit that we have uncovered may be accessed ‘emotionally,’ we envisage that the circuit may also be used in new ways to restore lost locomotor function e.g. in Parkinson’s disease or after spinal cord injury.

In summary, we reveal a crucial link between neural circuits and adaptive actions, demonstrating how internal states encoded in the hypothalamus significantly influence behavior. The LHA-PPN projections, which prioritize safety over other essential needs, challenge conventional notions of movement control. The findings demonstrate how brain circuits orchestrate context-dependent locomotion, bypassing cortical and basal ganglia inputs in favor of a direct excitatory projection from deep brain structures to the midbrain. As our comprehension of neural circuits expands, we uncover new avenues to comprehend how the brain orchestrates adaptive actions. Our findings resonate with the primal instinct to seek safety, a phenomenon that transcends species and underscores the delicate balance between survival and exploration.

## Acknowledgements

We thank the Clemmensen and Kiehn lab members for their discussions and input to the study. Special thanks to Debora Masini for her assistance in the Ethovision protocol experimental setup. We also extend our gratitude to Jared Cregg and Michael Andersen for their help with fiber photometry setup. Rhagavendra Selvan and Daniel Jercog provided valuable support in the scriptwriting for the fiber photometry analysis. We thank Iryna VesthHansen for genotyping the Vglu2-cre colony. This research is supported through the International Postdoc Fellowship of the Novo Nordisk Foundation Center for Basic Metabolic Research (NK), Lundbeck foundation (OK: R345-2020-1769) and Novo Nordisk Foundation Laureate Program and its Continuation (OK: NNF15OC0014186 /NNF24SA0088394), the Novo Nordisk Foundation (CC: NNF22OC0073778) and grants to the Center for Basic Metabolic Research (NNF18CC0034900 and NNF23SA0084103).

## Author contributions

C.C., O.K. and N.K. conceptualized the study, Methodology, C.C., O.K. and N.K.; N.K. performed technical procedures for the anterograde and retrograde tracing, optogenetic activation, chemogenetic activation and fiber photometry studies and performed the respective data analysis. L.S. performed surgeries and behavioral studies, the chemogenetic inhibition and anatomical tracing and evaluation; C.C O.K. and N.K. wrote the original draft; All authors reviewed and edited the manuscript. O.K and C.C. supervised all aspects of the study and acquired funding.

## Methods

### Experimental animals

#### Mice

Animal procedures were performed in accordance with European Union Directive 2010/63/EU, and approved by the Danish Animal Inspectorate (Dyreforsøgstilsynet, permit 2017-15-0201-01172 and 2022-15-0201-01286) as well as the clinical veterinarians at the Department of Experimental Medicine, Faculty of Health and Medical Sciences, University of Copenhagen (plans P21-297, P22-549, A20-160 and A23-154). Experiments were performed in adult mice of both genders and older than 7-8 weeks. Heterozygous Vglut2Cre mice55 and VGATCre (Jackson 016962) were used for experiments. Mice were housed in a group setting in ventilated cages at 23-24 °C and 45-50% humidity. The mice were maintained on a 12-hour light and dark cycle and were provided with ad libitum access to food and water, except during fasting periods.

### Surgical procedures

#### Stereotaxic injections

Surgery was performed in a stereotaxic setup (Stereotaxic frame Model 900, David Kopf) coupled to the StereoDrive software (Neurostar. Mice received subcutaneous injections of Buprenorphin (0.1 mg/kg of body weight, 472318, Salfarm Danmark A/S) and a local injection of lidocaine (550147, Salfarm Danmark A/S) prior to the surgery. Viscotears (587852, Alcon) was used to protect the eyes during the surgery. Mice induced with 5% isoflurane (055226, ScanVet) and maintained on a constant flow of oxygen enriched 1.5-2% isoflurane. Mice were kept at a stable temperature during the procedure by using a heating pad (53800M, Stoelting). Fine scissors were used to open the scalp, and the exposed skull was cleaned and dried before the set up was calibrated to tilt and scale using Bregma, Lambda and landmarks of 2mm left/right of midline. After determining the target location, a small hole was drilled into the skull using a handheld drill (Dremel). Postsurgical treatment after viral infection and probe implantations included a subcutaneous injection of Metacam (2 mg/ml, QM01AC06, Boehringer Ingelheim) to relieve pain and inflammation. In the days following the surgery, mice were examined daily for discomfort and received Buprenorphine in a nut cream (0.08 mg/g) for pain management in the amount that corresponded to the subcutaneous injection dose.

##### Viral infection

Viral suspensions were injected using a fine glass micropipette (Neurostar). Within the brain area, the needle was moved at a speed of 0.1 mm/s to ensure minimal invasion. For accurate injection volume a Neurostar injectomate was used at a rate of 50 nl/min with a delay of 6 min after injection to extract the needle. After injection, the skin covering the skull was closed with small wound clips (203-1000, Agnthos) or sutures (114026, Ethicon).

##### Probe implantation

Optical fibers were mounted utilizing a cannula holder (SCH_1.25, Doric Lenses). Probes were lowered into the brain at a speed of 0.1 mm/s to ensure minimal invasion. When the target region was reached, the probe was fixed in place with UV hardening dental (VI64413B, Ivoclar Vivadent) cement and two component adhesives (36658, Optibond). To harden the cement and adhesive a handheld UV lamp (100045, M+W Dental) was used.

#### Stereotaxic coordinates for viral injections and fiber implantation

**Pedunculopontine Nucleus (PPN):** 80 nl of viral suspension was injected at a sagittal angle of -20º with anterior-posterior (AP) -4.7 mm, mediolateral (ML) -1.28 mm, and dorsal-ventral (DV) -3.75 mm coordinates measured from Bregma. Fiber implants for optogenetics [0.22 numerical aperture (NA), 200 µm core diameter] were placed above the PPN (−20º sag. angle; AP -4,84 mm, ML -1.30 mm and DV -2.99 mm). **Lateral hypothalamus (LHA):** 150 nl of viral suspension was injected at (from bregma): AP -1.6 mm; ML -1.0 mm; and DV -5.0 mm); fiber implants for optogenetics [0.22 numerical aperture (NA), 200 µm core diameter] were placed above the LHA at AP -1.6 mm, ML -1.0 mm, and DV -4.6 mm. Fiber implants for fiber photometry were placed in the LHA at AP -1.6 mm, ML mm -0.88, and DV -4.9 mm. **Ventral tegmental area (VTA):** 80 nl of viral suspension was injected at (from bregma): AP -3.0 mm, ML -0.55 mm, and DV -4.45 mm. **Locus coeruleus (LC):** 80 nl of viral suspension was injected (from bregma): AP -5.5 mm, ML 0.9 mm, DV -3.70 mm). **Lateral habenula (LHb):** 80 nl of viral suspension was injected at (from bregma): AP-1.58 mm, ML -0.45 mm, and DV -2.84 mm). Stereotaxic coordinates for brain regions were taken from ‘The mouse brain in stereotaxic coordinates’.^56^

#### Viral vectors in experiments

For retrograde anatomical tracing experiments in Figure 1 and 1S: AAVretro-hSyn1-dlox-H2bj-mScarlet-I-NLS-EGFP(rev)-dlox-WPRESV40pA (VVF Zurich; v588-retro; 6.6×10e12) was injected into PPN. For anterograde anatomical tracing experiment in Figure 1 and 1S: AAV1-phSyn1(S)-FLEX-tdTomato-T2A-SypEGFP-WPRE (Salk Institute, 2.33×10e12) was injected in LHA. For optogenetic activation of glutamatergic LHA-PPN projection neurons in Figures 2, 2S, 3, 3S, 5, 5S: AAVretro-EF1a-doublefloxed-hChR2(H134R)-EYFP-WPRE-HGHpA (Addgene; 20298-AAVrg; 1×10e13) was injected into PPN. For optogentic activation of the glutamatergic PPN in Figure 3S: AAV5-Ef1a-DIO-hChR2-(E123T/T159C)-eYFP-WPRE was injected into the PPN (Addgene, 35509-AAV5, 4×10e12). For chemogenetic experiments in Figure 4 and 4S: AAVretro-FLEX-EGFP-2A-FlpO (VVF Zurich; v171-retro; 7.3×10e13) was injected into PPN. The second virus injected for activation AAV5-hSyn1-dFRT-hM3D(Gq)-mCherry-WPRE (VVF Zurich; v189-5; 3.5×10e12) and the second for inhibition AAV8-hSyn1-dFRT-hM4Di-mCherry-dFRT-WPRE-hGHp(A) (VVF Zurich; V190-8; 5.2 x10e12) were injected into LHA. For fiber photometry experiments Fig 4 and 4S: dual virus strategy of AAVretro-pEF1a-DIO-FLPo-WPRE-hGHpA (Addgene; 87306-AAVrg; 7×10e12) in PPN and AAVDJ-hEF1a-dFRT-jGCaMP7s(rev)-dFRT-WPRE-hGHp(A) (VVF Zurich; v521-DJ; 7×10e12) in LHA. For collateral tracing in Figure 5 and 5S: ssAAV-retro/2-hEF1α-dlox-EYFP(rev)-dlox-WPRE-hGHp(A) (VVF Zurich, V343-retro, 6.2×10e12) was injected into PPN and AAVretro-hSyn1-FLEX-mCherry (VVF Zurich, v116-retro, 4.8×10e12) was injected into VTA, LHb or LC respectively. For chemogenetic inhibiton of PPN or VTA in combination with optogenetics in Figure 6 and 6S: ssAAV-5/2-hSyn1-dlox-hM4D(Gi)_mCherry(rev)-dlox-WPRE-hGHp(A) (VVF Zurich; V84-5; 5.6×10e12) was injected in PPN and VTA respectively.

### Optogenetic experiments

#### Stimulation parameters

For all optogenetic experiments, pulse trains were generated using a Master 9 pulse stimulator (A.M.P.I.). Light was generated by Laserglow (blue light; 473 nm) and Optoduet (yellow light; 593 nm; IkeCool) lasers. Triggers were managed through Ethovision software protocols (Noldus Information Technology) using TTL pulses through a mini-USB-10 box (Noldus Information technologies).

Trains of stimuli varied between 10-80 Hz with a fixed pulse duration of 10 ms. For general optogenetic activation protocols a blue light stimulation with 473 nm a frequency of 40 Hz and pulse duration of 10 ms was used. Laser power settings were adjusted for each mouse and maintained throughout all experiments. The power of the laser was between 9-20 mW at the connector. Control stimulation with yellow light (593 nm) used 40 Hz, 10 ms pulse duration and peak light power 40 mW at the connector. For optogenetic inhibition a constant light stimulation was used for the length stated in the description of the respective experiment. For control stimulation we used the same duration of constant light at 593 nm, and peak light power of 2–3.5 mW at the connector.

#### Behavioral tests

##### Frequency testing for optimal locomotor performance in LHA

To identify the frequency optimal to evoking movement from LHA stimulation, *Vglut2*^*Cre*^ virus injected, and probe implanted mice were placed in the middle of an empty arena (50 cm x 50 cm) for free exploration. The laser was triggered randomly at 10, 20, 40, 50, and 80 Hz (stimulation lasted 3 seconds with 10 ms pulse duration). Pre-laser, laser-on, and post-laser periods were each set to three seconds. The distance moved and velocity were recorded using Ethovision software (Noldus Information Technology). After exporting raw data from Ethovision, the results were represented in graphs utilizing Prism GraphPad.

##### Real-time place preference optogenetics

Virus injected and probe implanted *Vglut2*^*Cre*^ mice were placed into a 50 cm x 50 cm arena that was segregated into two zones (each 25 cm x 50 cm) by walls with an open entrance zone (10 cm) between the zones. During the initial 10-minute session, the laser was activated whenever the mouse remained in zone A for at least two seconds. During the second half of the 10-minute trial, the mouse received laser stimulation when situated in zone B for at least 2 seconds. The laser stimulation was 40 Hz for 1 second. Mice were always placed in zone A at the beginning of the experiment. Behavioral parameters were analyzed using Ethovision software. Time in zone and heat maps were analyzed using arena specifications and time slots. Data were plotted and statistically analyzed using Prism GraphPad. Heatmaps were plotted using Ethovision.

##### Shelter and social zone optogenetics

To determine whether optogenetic activation of the brain area of interest could induce a preference for seeking shelter over seeking social interaction we performed social interaction experiments on virus injected and probe implanted *Vglut2*^*Cre*^ *and* control mice. The experimental arena measured 50 cm x 50 cm and included a custom-made shelter (3D printed on PRUSA 3D printer, height: 9.3 cm; width: 11.5 cm; length: 8.5 cm) placed in one corner and a social interaction cage positioned in the opposite corner. A novel mouse of the opposite sex was introduced to the social interaction cage (height:16.5cm; diameter: 7cm). To prevent any bias towards a particular corner based on the behavior room’s layout, prior to the social mouse experiments, we rotated the arena, so that the social and shelter corners of the arena switched. The quarter of the arena that contained the social chamber was designated as the “social quarter”. The entire experiment lasted 12 minutes, with 2 minutes of habituation and 10 minutes of active experimentation. During each activation trial, the laser was triggered 1.5 seconds after the mice entered the social quarter. To prevent photodamage, laser stimulation with 473 nm or 593 nm was administered at 40 Hz for a duration of 1.5 seconds, with a maximum of 30 stimulations per exploration.

##### Shelter and food zone optogenetics

This test was used to explore whether LHA circuit activation will cause a shelter-seeking in competition with food seeking after fasting. The arena (50 cm x 50 cm) contained a shelter (h: 9.3 cm; w: 11.5 cm; l: 8.5 cm) in one corner and a petri dish (10 cm diameter) with food placed in the opposite corner. Virus injected and probe implanted mice were fasted for 16-18h prior to the experiment. The quarter of the arena with the food dish was designated as the ‘food quarter’. The total time of the experiment was 12 minutes, with 2 minutes of habituation and 10 minutes of active experiment. During the active experiment, whenever mice were in the food quarter for 1.5 seconds, the optical trigger was activated after another 1.5 seconds. The laser stimulation with 473 nm or 593 nm was performed at 40 Hz with a duration of 1.5 seconds, with a maximum 30 times per exploration.

#### Analysis of shelter-seeking

With the Ethovision software the times spent in zones were analyzed using arena designation and velocity was determined using the 1.5-second-long time slots for pre-laser, laser-on and post-laser periods. Kaplan-Meier survival plot data for optogenetic experiments were extracted from the raw data by determining time stamp of laser trigger start to mouse in shelter (detected in shelter entrance or not detected). All events of the mice out of the shelter were plotted together for the survival plot. Time back to shelter was plotted as survival (1) in the case of the mouse not returning before the end of the trial this time was plotted as death (0). Mice that remained in the shelter for the entire experiment were excluded from the analysis. “N” corresponds to the number of animals in each group and “n” corresponds to the number of triggers in total.

### Chemogenetic experiments

Clozapine-N-Oxide (CNO, 4936, Tocris) was dissolved in 0.9% saline (220864, Braun) at a concentration of 0.1 mg ml^−1^. Saline or CNO (1-3 mg/kg) and was administered intraperitoneally. To test the effectiveness of the solution, each newly prepared CNO solution was tested in a control animal before starting the experiment. To reduce a confounding factor of stress induced by handling and injection all experimental animals were sham treated by manually handling them daily for two weeks prior to the start of the experiments. To prevent any possible habituation effect, the study was designed as a crossover experiment, randomizing the animals into two groups. One group was initially administered saline and then CNO, while the other group received CNO first and then saline. To prevent animals from transferring any experimental stress to cage mates, we utilized single housing in cages before and after the experiment. To ensure full effectiveness, CNO was administered 20 minutes before the start of the experiment. As CNO can remain biologically effective for up to 72 hours,^33,34^ we conducted mouse testing with a minimum of 3-day pause to ensure the drug had washed out. To enhance the novelty of the experiment, the walls were patterned, and the arenas were repositioned within the room. In the first round of experiments, the arenas were cleaned with 70% ethanol (83801380-5LC10, VWR) between the mice, whereas a window cleaning solution was used for the crossover round of experiments. The same experimental arena was utilized, with a minimum gap of two weeks between each trial to avoid familiarity in the mice.

#### Chemogenetic behavioral tests

##### Open field arena

To determine whether chemogenetic stimulation affected general locomotor activity, viral injected mice were subjected to an open field test. Mice were injected with CNO or saline 20 minutes prior to placement into the middle of an open field arena (50 cm x 50 cm). Exploration was recorded for 15 minutes and analyzed with the Ethovision software.

##### Shelter and social zone

Twenty minutes after injection, the mice were placed in a 50 cm x 50 cm arena with a shelter (h: 9.3 cm, w:11.5 cm, l: 8.5 cm) in one corner and a social cage housing a novel mouse of the opposite gender. The social cage was placed in the opposite quarter of the shelter. The mice were recorded for a total of 12 minutes, with the first 2 minutes serving as a habituation period. Ethovision software was used to analyze videos, and exported raw data was utilized to analyze the time it took the mice to reach the shelter.

##### Shelter and food zone

Mice were fasted for 16 hours overnight, with free access to water. The next day, mice were injected with CNO or saline and 20 minutes later placed into the arena (50 cm x 50 cm) containing a shelter (h: 9.3 cm; w: 11.5 cm; l: 8.5 cm) in one corner and a petri dish with food pellets in the opposite corner. The trial duration was 12 minutes total, including 2 minutes of habituation. Trial videos were recorded and analyzed using Ethovision software. As in the social zone trial, time to shelter was measured based on the raw data from Ethovision.

##### Shelter and ultrasound

To test if cue directed shelter-seeking was affected by chemogenetic inactivation, we utilized an arena containing an ultrasound speaker (Copenhagen University Electronics Workshop). The arena (50 cm x 50 cm) contained a shelter (h: 9.3 cm; w: 11.5 cm; l: 8.5 cm) in one corner and a custom-made small speaker transmitting 20 kHz frequency when triggered in the opposite corner. Total trial time was 12 minutes, with 2-minute habituation. The ultrasound was turned on whenever the mouse entered the sound zone. Videos were recorded with Ethovision and time back to shelter was extracted from the Ethovision raw data export.

#### Analysis

The Kaplan-Meier survival plot data for chemogenetic experiments was extracted from the raw data by determining time stamps detected in the zones of interest (food, social or ultrasound) to mice in shelter. Data for the velocity curves was extracted from the Ethovision raw data, where each velocity data point corresponds to 0.08 seconds. In the case of not enough triggers in the shelter-seeking arena, the number of analyzed traces was adjusted to the compared track (i.e. 473 nm compared to 473 nm + CNO.) “N” corresponds to the number of animals in each group and “n” corresponds to the number of triggers in total.

#### Fiber photometry

### Recordings

Recordings were conducted using the TDT Lux RZ190X photometry setup (Tucker Davis Technologies), in conjunction with the Synapse software (Tucker Davis Technologies) and a USB camera (960-001055, Logitech). To ensure optimal signal quality, 5-millimeter-long fiber optic cannulas with metal ferrules (MFC_400/430-0.66_5mm_MF1.25_FLT, Doric Lenses) were implanted into the brain using the same procedure implemented for the optogenetic experiment’s fiber optics. The patch cord was connected to a rotary rod to improve signal quality. The LEDs’ power output was adjusted, resulting in 20 mW and 25 mW for the 405 nm and 465 nm channels, respectively, according to the TDT power meter. The TDT Synapse software aligned frames to the photometry signal and timestamped the video. A TTL (Transistor-Transistor Logic) connection enabled precise triggering of ultrasound and airpuff.

#### Behavioral tests

##### Spontaneous activity

Spontaneous calcium activity was tested in an open field (50 cm x50 cm) and mice with adequate signal were selected for behavioral experiments.

##### Arena with food zone and airpuff

Mice were placed in an arena with a food zone located in the opposite corner of the shelter (h: 9.3 cm; w: 11.5 cm; l: 8.5 cm). A tube connected to a picospritzer (Parker) released a 500 ms airpuff of 50 psi when triggered and was placed above the food zone. Each time the mouse approached the food zone, an airpuff was activated. The mice were recorded for 15 minutes.

##### Arena with food zone and airpuff sound

To control for neural activity during approach to the food zone and response to the sound of the airpuff, mice underwent recording in control experiments without airpuff. Here we used the same arena as in “food zone and airpuff”, with the difference that the tube was not connected to the picospritzer. When the mouse approached the food, the airpuff sound (500 ms) was triggered.

##### Ultrasound arena with shelter

Mice were recorded for 10 minutes in an arena that had a shelter (h: 9.3 cm; w: 11.5 cm; l: 8.5 cm) in one corner and a small ultrasound speaker with food in the opposite corner. When the mouse approached the speaker, a 20 kHz sound was triggered for 1.5 seconds.

#### Analysis

Fiber Photometry analysis was performed using custom-made python scripts. Artifacts were removed from the beginning of the recording by analyzing the sample starting at 10 seconds post-recording start time. Sampled traces were down sampled by a factor of 10. Traces of different animals were smoothed and detrended. The dF/F was determined through the demodulated signal. The z-scores for individual trials were calculated by subtracting the mean and dividing by the standard deviation from a reference period of -5 seconds to the onset of the stimulus. TTL input was used to identify the onset and offset of the airpuff and ultrasound. The determined onset times were utilized to generate z-score snippets for all subjects, which were subsequently employed to generate z-score graphs depicting ultrasound responses. Heatmaps were plotted with all cue events sorted by Area under the curve (AUC) score. AUC comparisons were determined from 1 second before and 1 second after the cue onset for the airpuff experiment. For the ultrasound experiment AUC was calculated 2 seconds before the cue onset and 2 seconds after, since the cue was 1.5 seconds long. All triggered cues were pooled together and compared. The script used was a modified version of the “Lick bout analysis” example on the TDT website. (TDT Offline Data Analysis tools: Lick Bout Epoc Filtering: https://www.tdt.com/docs/sdk/of-fline-data-analysis/offline-data-python/examples/LickBouts/).

### Anatomical evaluation of collaterals along LHA anterior posterior axis

Images were taken of slices throughout the extent of the anterior-posterior LHA axis using a Axio imager 2 microscope (Zeiss) and Axio Observer microscope (Zeiss) at 10x magnification. Images were processed using the Zen software (Zeiss). Cells located within the LHA were determined and classified as mCherry or GFP positive and cells expressing both mCherry and GFP were further determined using the Zen Lite software (Zeiss). All images were further processed using Adobe Illustrator (v.28) and cells determined as mCherry, GFP or mCherry/GFP positive counts. Only cells in the ipsilateral LHA of the viral injections were counted.

#### Post experimental analysis of brain tissue

All brains were analyzed post-experimentally to validate injection sites and retrograde labeling of cell bodies. For the perfusion, mice were sedated with pentobarbital (250 mg/kg Department of Experimental Medicine, University of Copenhagen) and transcardially perfused with fresh 50 ml cold PBS

(Phosphate Buffered Saline, 0.05 M) and heparin (10 U/ml, A90402, LEO Pharma) followed by 4% paraformaldehyde solution (HL96753.1000, Histolab). The brains were then isolated and post-fixed in 4% PFA for 4 hours. After post-fixing, brains were placed in a 25% sucrose solution overnight at 4 °C. Using a cryoprotectant solution (Neg-50, 6502, Epredia), brains were frozen on dry ice and kept at -80 °C until cryo-sectioning. Sections were sliced in a CryoStar NX70 cryostat (AxLab, Thermo Fisher Scientific) at 30 µm and collected on a glass slide.

#### Immunochemistry of brain tissue

To ensure coherence within results, all samples of one experiment were stained in the same round. After washing, the sections were blocked with 5% of the respective serums NGS normal goat serum (AB_2336990, Jackson Immunoresearch) or NDS normal donkey serum (AB_2337258, Jackson Immunoresearch) in PBS-T (0.5% Triton-X100, 9036-19-5 Sigma Aldrich, in PBS, 10010023 Gibco). For overnight staining at 4°C, the primary antibodies (anti-DsRed in rabbit 1:1000 (632496, Takara); anti-GFP in chicken 1:1000 (13970, Abcam) were applied in solution (PBS-T, 1% serum), together with the nuclear counterstaining NeuroTrace 435/455 (1:500, N21479, Thermo Fisher Scientific). After washing, the secondary antibodies coupled with fluorophores (1:500 Alexa-568 anti-rabbit, A10042, invitrogen; 1:500 Alexa488 anti-chicken, A11039 invitrogen) were applied in solution (PBS-T, 1% serum) and the sections were incubated for 2 hours at room temperature. Post-secondary staining, sections were washed in PBS and mounted on microscope slides (J1800AMNZ, Thermo Scientific) using Mowiol mounting medium (475904-M, Sigma Aldrich) and coverslips (BBAD02400600#SC13MNZ, Menzel Glaeser).

#### Imaging of brain slices

Brain slices were imaged as tile images to enable determination of the exact anatomical location of virus expression and fiber placement. Imaging of the tile images was done at a magnification of 10x with an epi-fluorescence microscope in the ZEN Pro Software (Zeiss). To verify exact labeling of cells and to show axonal terminals, imaging was performed using a confocal microscope. All brain slices within an experiment were imaged with the same settings to reduce variability.

### Statistical analysis and quantification

All statistical analyses were executed in Prism 9 (GraphPad). For comparison of two groups, a two-tailed Student t-tests was used. For comparisons of the same group in different settings, a paired t-test was used. In experiments with multiple groups, comparisons were performed using one-way ANOVA or two-way repeated measures ANOVA. Survival plots were compared using a Kaplan-Meier test (Log-rank test). Statistical representations are shown as mean ±SD with p-values below 0.05 being considered significant (^*^p< 0.05, ^**^p < 0.01, ^***^p < 0.001, ^****^p < 0.0001 and ns = not significant).

**Extended Data Figure 1:**
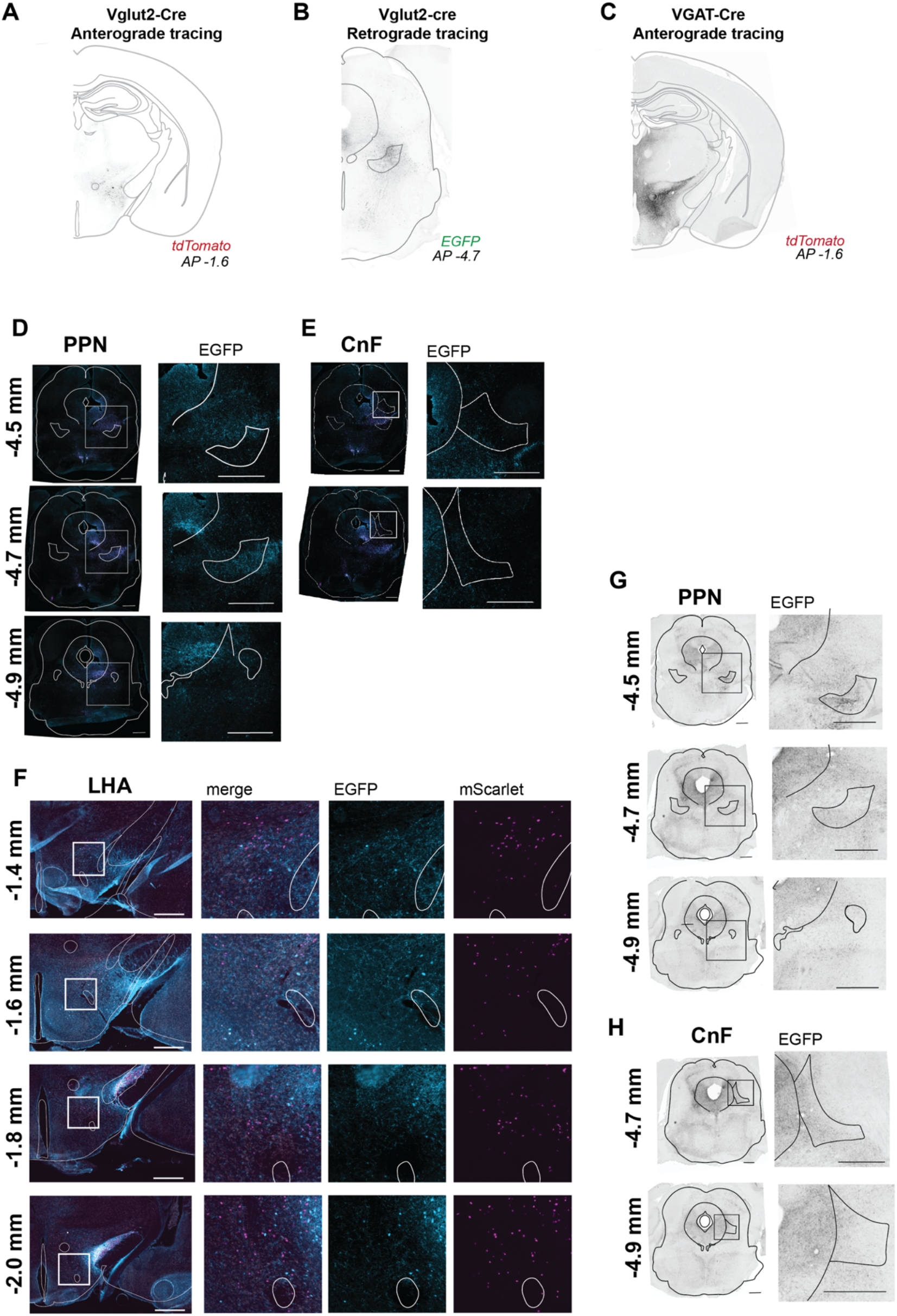
Retrograde and anterograde tracing reveal glutamatergic projections from Lateral Hypothalamic area (LHA) to the caudal pedunculopontine nucleus (PPN). **A)** Representative tile image of anterograde injection site in LHA with tdTomato expression in Vglut2^+^ cells (AP -1.6 from Bregma) in *Vglut2*^*Cre*^ *mice*. **B)** Representative tile image of retrograde tracing injection site in PPN with EGFP expression (AP - 4.7 from Bregma) in *Vglut2*^*Cre*^ mice. **C)** Representative tile image of anterograde injection site in LHA with tdTomato expression in VGAT^+^ cells (AP - 1.6 from Bregma) in *VGAT*^*Cre*^ mice. **D)** Representative merged image and zoomed in images at AP -4.5, -4.7 and -4.9 mm from Bregma showing Vglut2 terminals in PPN (SypEGFP expression in *Vglut2*^*Cre*^ mice). **E)** Representative merged tile image and zoomed in images at AP -4.7 and -4.9 mm from Bregma Vglut2 terminals (SypEGFP expression in *Vglut2*^*Cre*^ mice). Lack of labeling in CnF. **F)** Representative merged tile image and zoomed in images at AP -1.4, 1.6, 1.8, 2.0 mm from Bregma. Retrogradely labelled cell bodies in LHA in both merged and separate channels for mScarlet and EGFP expression in *Vglut2*^*Cre*^ mice. Vglut2 positive cells show EGFP. **G)** Representative tile image and zoomed in images at AP -4.5, -4.7 and -4.9 mm from Bregma. Anterograde labelled terminals in PPN show SypEGFP expression in *VGAT*^*Cre*^ mice. **H)** Representative tile image and zoomed in images at AP -4.7 and -4.9 mm from Bregma. Anterograde labelled terminals in CnF show SypEGFP expression in *VGAT*^*Cre*^ (All scale bars 500µm)

**Extended Data Figure 2:**
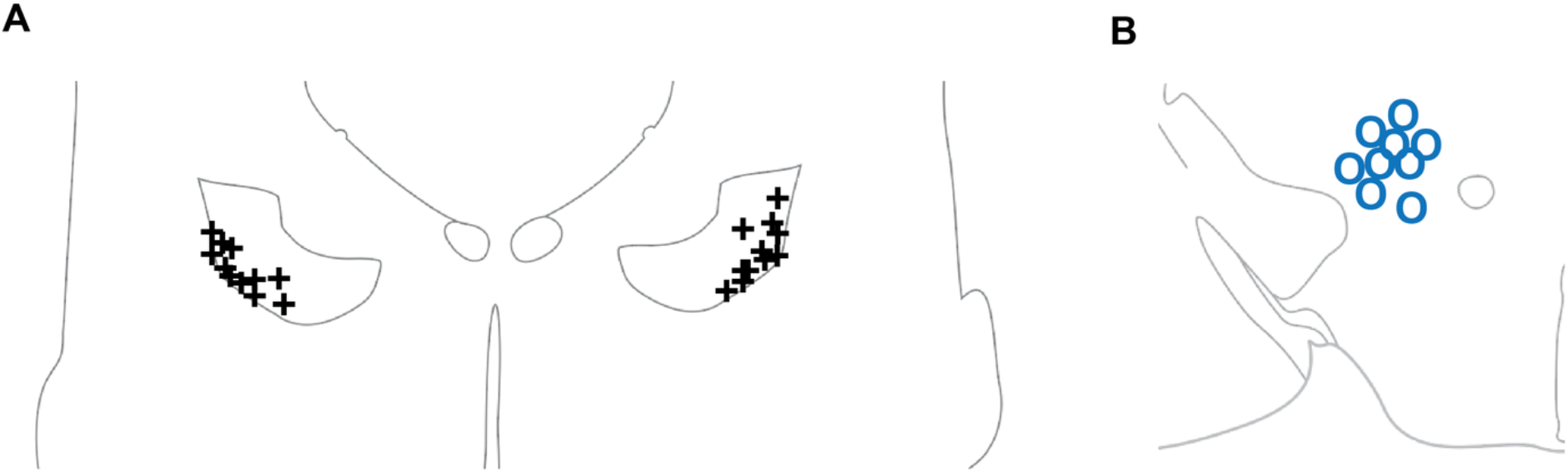
Viral injection sites in PPN and fiber placement sites in LHA. **A)** Estimated center of viral injection sites in PPN at -4.7 mm AP from bregma for all animals (80nl injected). **B)** Fiber placement sites in LHA at -1.6 mm AP from bregma for all animals.

**Extended Data Figure 3:**
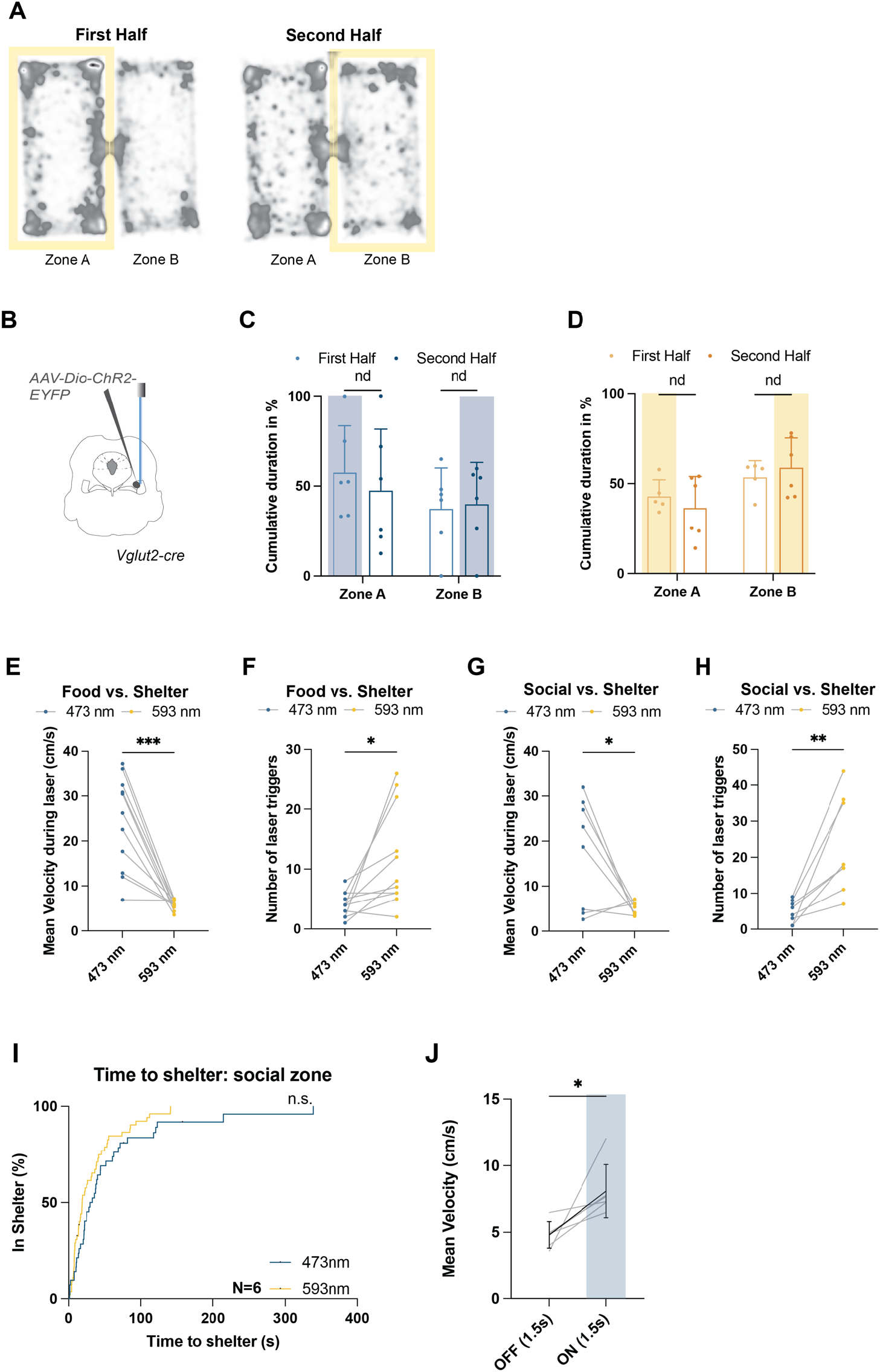
A. Yellow light stimulation of LHA-PPN projection neurons does not trigger place aversion. **A)** Heatmaps with traces of all control laser (593 nm) tested mice in the first half (0-10 mins, stimulation in Zone A) and second half (10-20 mins, stimulation in Zone B) of the trials. B-D. Broad optogenetic activation of Vglut2 positive neurons in the caudal PPN causes no place preference. **B)** Experimental strategy with injection of AAV5-Dio-ChR2-eYFP and fiber placement in PPN. **C)** Cumulative time (% of total time) in zones with 473 nm laser. (Zone A: Time spent first vs. second time period (N=6, mean ± SD, paired t-test p=0.311076); Zone B: Time spent first vs. second time period (paired t-test; p=0.530555). Blue highlighted area represents stimulated zone. **D)** Cumulative time (% of total time) in zones with 593 nm control laser. (Zone A: Time spent first vs. second time period (N=6, mean ± SD, paired t-test p=0.675364); Zone B: Time spent first vs. second time period (paired t-test; p=0.807286). Yellow highlighted area represents stimulated zone. E-H. Velocity of locomotion during shelter-seeking and number of stimulations in food or social zone following glutamatergic LHA-PPN projection neuron stimulation. **E)** Mean velocity of locomotion is higher during stimulation with the 493 nm than with the control 593 nm laser in the food vs. shelter arena (N=11; paired t-test p=0.0002; each data point represents one animal). **F)** Number of stimulations in food zone. (N=11; paired t-test p=0.0119; each data point represents one animal). **G)** Mean velocity of locomotion during stimulation with 493 nm was significantly higher than with 593 nm laser stimulation in social vs. shelter arena (N=8; paired t-test p=0.0258; each data point represents one animal). **H)** Number of stimulations in social zone. (N=8; paired t-test p=0.0044; each data point represents one animal). **I-J. Broad optogenetic activation of Vglut2 positive neurons in the caudal PPN causes no shelter-seeking. I)** Survival plot of latency of return time to shelter after laser stimulation shows no difference in comparison of 493 nm laser and 593 nm control laser (N=6; Kaplan-Meier (Log-Rank Test p=0.0708). **J)** Comparison of mean velocity before and after 473nm stimulation in the social vs. shelter arena shows increased velocity. Highlighted area shows laser stimulation; mean ± SD with single values in grey (N=6; paired t-test; p=0.0310).

**Extended Data Figure 4:**
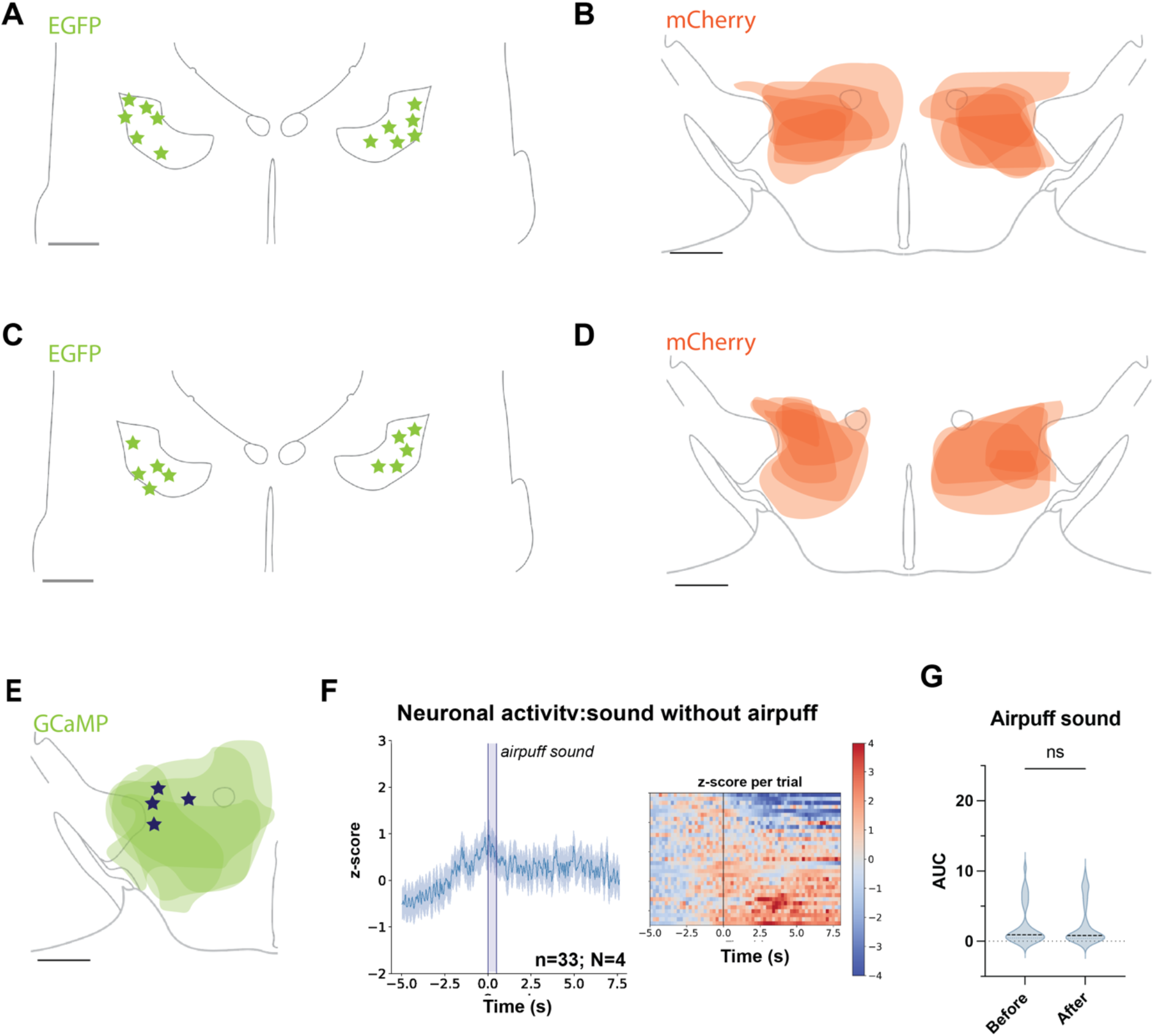
A-B. Virus injection site in caudal PPN and area for recombination in LHA for chemogenetic activation experiments. **A)** Estimation of center of *AAVretro-FLEX-EGFP-2A-FlpO* injection sites in PPN (EGFP, AP -4.7 from Bregma, scale bar 500 mm; N=6) in chemogenetic activation subjects. **B)** Virus infection spread in LHA (mCherry, AP 1.6 from Bregma, scale 500 mm): Expression of *AAV5-hSyn1-dFRT-hM3D(Gq)-mCherry-WPRE* in all subjects of the chemogenetic activation experiment (Opacity 30%, N=6). **C-D. Virus injection site in caudal PPN and area for recombination in LHA for chemogenetic inactivation experiments. C)** Estimation of center of *AAVretro-FLEX-EGFP-2A-FlpO* injection sites in PPN (EGFP, AP -4.7 from Bregma, scale bar 500 mm; N=5) in chemogenetic inactivation subjects. **D)** Virus infection spread in LHA (mCherry, AP 1.6 from Bregma, scale 500 mm): Expression of *AAV8-hSyn1-dFRT-hM4Di-mCherry-dFRT-WPRE-hGHp(A)* in all subjects of the chemogenetic inactivation experiment (Opacity 30%; N=5). E. GCaMP7 expression in LHA. **E)** Expression of *AAVDJ-hEF1a-dFRT-jGCaMP7s(rev)-dFRT-WPRE-hGHp(A)* in all subjects marked in green (30% opacity) with blue stars indicating fiber placement in LHA (AP 1.6 from Bregma, scale 500 um; N=4). F-G. LHA-PPN projection neuron activity after sound but no airpuff. **F)** Z-score plot of photometry signal after airpuff sound alone (Mean +/-SD; n=33 sound triggers of 500 ms, N=4 mice) and heatmap plot of z-scores for every trial separately. G) Area under the curve measurement for 1 second before and after sound only onset (Wilcoxon matched pairs signed rank test p=0.2798).

**Extended Data Figure 5:**
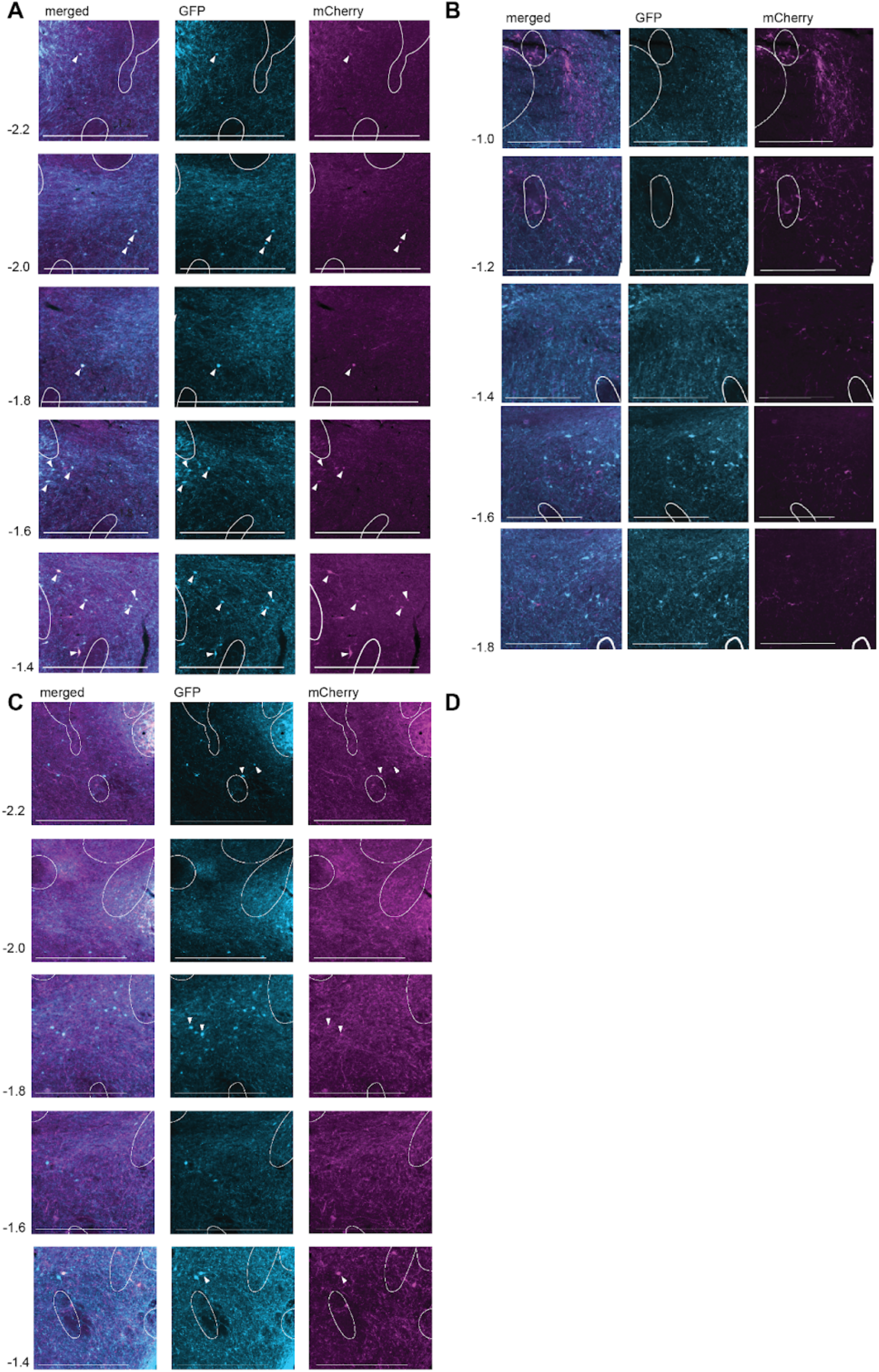
LHA-PPN-, LHA-VTA-, LHA-LHb, and LHA-LC projection neurons along the anterior-posterior (AP) axis of the LHA. **A)** Representative images of cell bodies projecting to PPN (*GFP*) and VTA (*mCherry*) along the LHA AP axis (numbers represent AP coordinate from bregma; scale bar 500 mm; double-labeled cells marked with white arrows). **B)** LHb (*mCherry*) and PPN (*GFP*) retrogradely labeled cell bodies along the AP axis of the LHA shown in separate channels or merged (numbers represent AP coordinate from bregma; scale bar 500 mm; double-labeled cells marked with white arrows). **C)** Representative images of retrogradely labeled cells from PPN (*GFP*) and LC (*mCherry*) along the LHA AP axis (numbers represent AP coordinate from bregma; scale bar 500µm, double-labeled cells marked with white arrows).

## References

1. Roseberry, T. K. et al. Cell-Type-Specific Control of Brainstem Locomotor Circuits by Basal Ganglia. Cell 164, 526–537 (2016).

2. Caggiano, V. et al. Midbrain circuits that set locomotor speed and gait selection. Nature 553, 455–460 (2018).

3. Josset, N. et al. Distinct Contributions of Mesencephalic Locomotor Region Nuclei to Locomotor Control in the Freely Behaving Mouse. Current Biology 28, 884-901.e3 (2018).

4. Masini, D. & Kiehn, O. Targeted activation of midbrain neurons restores locomotor function in mouse models of parkinsonism. Nat Commun 13, 504 (2022).

5. Arber, S. & Costa, R. M. Networking brainstem and basal ganglia circuits for movement. Nat Rev Neurosci 23, 342–360 (2022).

6. Branco, T. & Redgrave, P. The Neural Basis of Escape Behavior in Vertebrates. Annu. Rev. Neurosci. 43, 417–439 (2020).

7. Kim, L. H. et al. Integration of Descending Command Systems for the Generation of Context-Specific Locomotor Behaviors. Front Neurosci 11, 581 (2017).

8. Leiras, R., Cregg, J. M. & Kiehn, O. Brainstem Circuits for Locomotion. Annu. Rev. Neurosci. 45, 63–85 (2022).

9. Noga, B. R. & Whelan, P. J. The Mesencephalic Locomotor Region: Beyond Locomotor Control. Front. Neural Circuits 16, (2022).

10. Pernía-Andrade, A. J., Wenger, N., Esposito, M. S. & Tovote, P. Circuits for State-Dependent Modulation of Locomotion. Front. Hum. Neurosci. 15, (2021).

11. Ryczko, D. & Dubuc, R. Dopamine control of downstream motor centers. Current Opinion in Neurobiology 83, 102785 (2023).

12. Korotkova, T. & Ponomarenko, A. To eat? To sleep? To run? Coordination of innate behaviors by lateral hypothalamus. e-Neuroforum 23, 45–55 (2017).

13. Sinnamon, H. M. Preoptic and hypothalamic neurons and the initiation of locomotion in the anesthetized rat. Progress in Neurobiology 41, 323–344 (1993).

14. Sinnamon, H. M., Karvosky, M. E. & Ilch, C. P. Locomotion and head scanning initiated by hypothalamic stimulation are inversely related. Behavioural Brain Research 99, 219–229 (1999).

15. Marciello, M. & Sinnamon, H. M. Locomotor stepping initiated by glutamate injections into the hypothalamus of the anesthetized rat. Behavioral Neuroscience 104, 980–990 (1990).

16. Jordan, L. M. Initiation of Locomotion in Mammals. Annals of the New York Academy of Sciences 860, 83–93 (1998).

17. Mickelsen, L. E. et al. Single-cell transcriptomic analysis of the lateral hypothalamic area reveals molecularly distinct populations of inhibitory and excitatory neurons. Nat Neurosci 22, 642–656 (2019).

18. Stuber, G. D. & Wise, R. A. Lateral hypothalamic circuits for feeding and reward. Nat Neurosci 19, 198–205 (2016).

19. Stuber, G. D. Neurocircuits for motivation. Science 382, 394–398 (2023).

20. Oh, S. W. et al. A mesoscale connectome of the mouse brain. Nature 508, 207–214 (2014).

21. Goñi-Erro, H., Selvan, R., Caggiano, V., Leiras, R. & Kiehn, O. Pedunculopontine Chx10+ neurons control global motor arrest in mice. Nat Neurosci 26, 1516–1528 (2023).

22. Chen, J. et al. A Vagal-NTS Neural Pathway that Stimulates Feeding. Current Biology 30, 3986-3998.e5 (2020).

23. Nieh, E. H. et al. Inhibitory Input from the Lateral Hypothalamus to the Ventral Tegmental Area Disinhibits Dopamine Neurons and Promotes Behavioral Activation. Neuron 90, 1286–1298 (2016).

24. Stamatakis, A. M. et al. Lateral Hypothalamic Area Glutamatergic Neurons and Their Projections to the Lateral Habenula Regulate Feeding and Reward. J. Neurosci. 36, 302–311 (2016).

25. Jennings, J. H., Rizzi, G., Stamatakis, A. M., Ung, R. L. & Stuber, G. D. The Inhibitory Circuit Architecture of the Lateral Hypothalamus Orchestrates Feeding. Science 341, 1517–1521 (2013).

26. Jennings, J. H. et al. Visualizing Hypothalamic Network Dynamics for Appetitive and Consummatory Behaviors. Cell 160, 516–527 (2015).

27. Verma, D. et al. Hunger Promotes Fear Extinction by Activation of an Amygdala Microcircuit. Neuropsychopharmacol 41, 431–439 (2016).

28. Tseng, Y.-T., Schaefke, B., Wei, P. & Wang, L. Defensive responses: behaviour, the brain and the body. Nat. Rev. Neurosci. 24, 655–671 (2023).

29. Jensen, M. et al. Anxiolytic-Like Effects of Increased Ghrelin Receptor Signaling in the Amygdala. IJNPPY 19, pyv123 (2016).

30. Tóth, K., László, K., Lukács, E. & Lénárd, L. Intraamygdaloid microinjection of acylated-ghrelin influences passive avoidance learning. Behavioural Brain Research 202, 308–311 (2009).

31. Pentkowski, N. S., Litvin, Y., Blanchard, D. C. & Blanchard, R. J. Effects of estrus cycle stage on defensive behavior in female Long-Evans hooded rats. Physiology & Behavior 194, 41–47 (2018).

32. Singh, D. K., Hari Dass, S. A., Abdulai-Saiku, S. & Vyas, A. Testosterone Acts Within the Medial Amygdala of Rats to Reduce Innate Fear to Predator Odor Akin to the Effects of Toxoplasma gondii Infection. Front. Psychiatry 11, (2020).

33. Roth, B. L. DREADDs for Neuroscientists. Neuron 89, 683–694 (2016).

34. Sternson, S. M. & Roth, B. L. Chemogenetic Tools to Interrogate Brain Functions. Annu. Rev. Neurosci. 37, 387–407 (2014).

35. Mongeau, R., Miller, G. A., Chiang, E. & Anderson, D. J. Neural Correlates of Competing Fear Behaviors Evoked by an Innately Aversive Stimulus. J Neurosci 23, 3855–3868 (2003).

36. Rossi, M. A. et al. Transcriptional and functional divergence in lateral hypothalamic glutamate neurons projecting to the lateral habenula and ventral tegmental area. Neuron 109, 3823-3837.e6 (2021).

37. Schwarz, L. A. et al. Viral-genetic tracing of the input–output organization of a central noradrenaline circuit. Nature 524, 88–92 (2015).

38. Soya, S. et al. Orexin modulates behavioral fear expression through the locus coeruleus. Nat Commun 8, 1606 (2017).

39. Barbano, M. F. et al. Lateral hypothalamic glutamatergic inputs to VTA glutamatergic neurons mediate prioritization of innate defensive behavior over feeding. Nat Commun 15, 403 (2024).

40. Barbano, M. F. et al. VTA Glutamatergic Neurons Mediate Innate Defensive Behaviors. Neuron 107, 368-382.e8 (2020).

41. Woolf, N. J. & Butcher, L. L. Cholinergic systems in the rat brain: III. Projections from the pontomesencephalic tegmentum to the thalamus, tectum, basal ganglia, and basal forebrain. Brain Research Bulletin 16, 603–637 (1986).

42. Hallanger, A. E., Levey, A. I., Lee, H. J., Rye, D. B. & Wainer, B. H. The origins of cholinergic and other subcortical afferents to the thalamus in the rat. Journal of Comparative Neurology 262, 105–124 (1987).

43. Ford, B., Holmes, C. J., Mainville, L. & Jones, B. E. GABAergic neurons in the rat pontomesencephalic tegmentum: Codistribution with cholinergic and other tegmental neurons projecting to the posterior lateral hypothalamus. Journal of Comparative Neurology 363, 177–196 (1995).

44. Capelli, P., Pivetta, C., Soledad Esposito, M. & Arber, S. Locomotor speed control circuits in the caudal brainstem. Nature 551, 373–377 (2017).

45. Isa, T., Marquez-Legorreta, E., Grillner, S. & Scott, E. K. The tectum/superior colliculus as the vertebrate solution for spatial sensory integration and action. Current Biology 31, R741–R762 (2021).

46. Li, Y. et al. Hypothalamic Circuits for Predation and Evasion. Neuron 97, 911-924.e5 (2018).

47. Lecca, S. et al. Aversive stimuli drive hypothalamus-to-habenula excitation to promote escape behavior. eLife 6, e30697 (2017).

48. Sciolino, N. R. et al. Natural locus coeruleus dynamics during feeding. Science Advances 8, eabn9134 (2022).

49. Barcomb, K., Olah, S. S., Kennedy, M. J. & Ford, C. P. Properties and modulation of excitatory inputs to the locus coeruleus. The Journal of Physiology 600, 4897–4916 (2022).

50. Peyron, C. et al. Neurons Containing Hypocretin (Orexin) Project to Multiple Neuronal Systems. J. Neurosci. 18, 9996–10015 (1998).

51. Poe, G. R. et al. Locus coeruleus: a new look at the blue spot. Nat Rev Neurosci 21, 644–659 (2020).

52. Maness, E. B. et al. Role of the locus coeruleus and basal forebrain in arousal and attention. Brain Res Bull 188, 47–58 (2022).

53. Nieh, E. H. et al. Decoding Neural Circuits that Control Compulsive Sucrose Seeking. Cell 160, 528–541 (2015).

54. de Jong, J. W. et al. A Neural Circuit Mechanism for Encoding Aversive Stimuli in the Mesolimbic Dopamine System. Neuron 101, 133-151.e7 (2019).

55. Borgius, L., Restrepo, C. E., Leao, R. N., Saleh, N. & Kiehn, O. A transgenic mouse line for molecular genetic analysis of excitatory glutamatergic neurons. Molecular and Cellular Neuroscience 45, 245–257 (2010).

56. Franklin, K. B. J. & Paxinos, G. The Mouse Brain in Stereotaxic Coordinates. (Elsevier Academic Press, Amsterdam Heidelberg, 2008).

